# Simple Methods to Acutely Measure Multiple Timing Metrics among Sexual Repertoire of Male *Drosophila*

**DOI:** 10.1101/2025.11.11.687834

**Authors:** Yutong Song, Hongyu Miao, Dongyu Sun, Xiao Liu, Fan Jiang, Xuejiao Yang, Woo Jae Kim

## Abstract

Male *Drosophila* courtship behavior is a key model for studying temporal decision-making. While courtship index (CI) is widely used to quantify mating activity, other timing metrics like courtship latency, copulation latency (CL), and mating duration (MD) remain understudied. Traditional methods for quantifying these behaviors are often labor-intensive and prone to human error.

In this study, we present a protocol combining a modular chamber system and automated software (DrosoMating) to quantify 6 key timing metrics during male courtship. Our image-based video analysis enables precise identification of courtship and copulation events, as well as quantification of their timing and duration under controlled conditions. Validation shows <0.05% error rates and 98–99% agreement with manual scoring for CL, CI, and MD. The protocol supports genetic and neural circuit manipulations, detecting subtle genetic, social, and environmental behavioral variations.

By minimizing manual effort and standardizing data collection, this approach facilitates scalable, reproducible studies on adaptive trade-offs, learning, and neural mechanisms in mating behavior. This method streamlines timing analysis in male courtship, offering reproducible metrics for behavioral genetics.

**Highlight:** A high-throughput software pipeline for automated temporal profiling of Drosophila melanogaster mating behavior after brief user-guided calibration.

An accompanying, open-hardware platform that can be assembled at minimal cost while maintaining experimental rigor.

The system attains near-manual accuracy and outputs Temporal Measurement Parameters data that are readily adaptable to—and quantifiable within—diverse behavioral paradigms.

## Introduction

Time perception is critical for survival, enabling organisms to estimate intervals between events and execute adaptive behaviors. Male *Drosophila* courtship behavior, a genetically well-characterized model (Singh and Singh, 2016), provides a robust framework for studying temporal control in action sequences. A widely used metric to quantify mating activity is the ‘Courtship Index (CI)’, defined as the percentage of time a male engages in courtship behaviors (e.g., following, wing vibration, abdominal curling) toward a female during a standardized observation window. CI reflects both the intensity and persistence of courtship, serving as a sensitive readout for dissecting genetic, environmental, or experimental perturbations on mating behavior, as demonstrated in studies investigating genetic mutations, environmental perturbations, and neuronal manipulations (Benton, 2015; Greenspan and Ferveur, 2000a; Kim, 2009; Pavlou and Goodwin, 2013; Yamamoto and Koganezawa, 2013).

Despite extensive research on courtship displays (Chen et al., 2024a), other critical metrics remain poorly characterized, especially as their representative characteristics as timing behaviors. ‘Courtship latency’ (time until a male initiates courtship after detecting a female) may reflect motivational state or sensory processing deficits but is rarely quantified in terms of genetics (Eastwood and Burnet, 1977a). ‘Copulation Latency (CL)’ (time from courtship initiation to successful mating) and ‘Mating Duration (MD)’ (length of mating itself) are similarly underexplored, though both likely impact reproductive success. Prolonged MD enhances paternity assurance by maximizing sperm transfer and suppressing female remating, but it simultaneously increases predation risks and energy expenditure (Bretman et al., 2009). Conversely, reduced MD may decrease a male’s paternity share due to incomplete sperm transfer or female resistance, yet it reduces exposure to ecological threats (e.g., predators, rivals) and conserves metabolic resources (Lee et al., 2023a). These traits may be influenced by cryptic factors such as male age, rival presence, or environmental stressors (e.g., temperature), yet their genetic and neural bases are unclear. While studies focus on stereotyped courtship actions, these “temporal metrics” offer insights into decision-making, persistence, and adaptive trade-offs—areas ripe for investigation (Huang et al., 2024; Kim et al., 2012a, 2013a, 2016, 2024; Lee et al., 2022; D. Sun et al., 2024; Y. Sun et al., 2024; Wong et al., 2019; T. Zhang et al., 2024b, 2024a; X. Zhang et al., 2024).

Numerous software tools have been developed to measure male mating behavior in *Drosophila*, yet many laboratories still rely on manual methods. Traditional approaches, such as the “courtship wheel”, have been widely used to assess male courtship activity (Hall, 1994; Neckameyer and Bhatt, 2016; Vosshall, 2007). However, quantifying the full repertoire of mating behavior metrics has historically been labor-intensive (Ejima and Griffith, 2007; Sokolowski, 2001). Recent advances in automation have addressed these challenges. For instance, Dankert et al. (2009) introduced a machine vision system to quantify male-female interactions, though its application has been more focused on studying male-male aggression (Dankert et al., 2009a). With the rapid development of machine learning and image processing, tools such as ‘Ctrax’ (Branson et al., 2009), ‘idTracker’ (Pérez-Escudero et al., 2014), Cadabra (Stern et al., 2015), ‘FlyTracker’ and MATLAB combined with ‘JAABA’ (Barwell et al., 2021; Kabra et al., 2013) have enabled precise quantification of male-female mating behavior. Recently, Chen et al. (2024) introduced an image-processing-based system that addresses this gap by automatically recognizing multiple courtship elements with high accuracy (>97%) and robust identity tracking even during complete fly overlap(Chen et al., 2024b). Despite these advancements, recent efforts to simplify courtship conditioning protocols (Gil-Martí et al., 2023a) still lack precise quantification for temporal parameters of post-copulatory behavior. Consequently, there is a need for more streamlined protocols and software to accurately measure the timing aspects of male mating behavior in *Drosophila*.

Beyond *Drosophila*-specific tools, general behavioral video analysis paradigms have been developed to track the movement and behavior of various organisms, including fruit flies. These methods often focus on tracking the position of single or multiple flies (Iyengar et al., 2012; Qu et al., 2022; Simon et al., 2011) or using artificial markers (Gal et al., 2020). Additionally, programs like ‘ToxTrac’ (Rodriguez et al., 2017), ‘UMATracker’ (Yamanaka and Takeuchi, 2018), ‘FIMTrack’ (Risse et al., 2017, 2014), and ‘ilastik’ (Berg et al., 2019) have been employed to analyze body posture, behavior, location, and movement trajectories in diverse species, including flies, bees, mice, and zebrafish. However, these tools often fall short in capturing the nuanced courtship behaviors of *Drosophila*, as motion data alone may not sufficiently represent complex interactions.

To address these limitations, we have developed a specialized software suite called ‘DrosoMating’ for analyzing and quantifying timing behaviors within male *Drosophila* courtship. By establishing 6 key indicators related to male courtship, we provide a robust framework for quantifying male mating behavior metrics under controlled conditions. This approach significantly reduces the time and errors associated with manual identification. Through iterative optimization, our system achieves an error rate of less than 0.05% compared to manual scoring. We believe this toolset offers a novel and efficient method for studying timing in male courtship behavior, providing valuable data and theoretical insights while streamlining the analysis process.

## RESULTS

### Video-Processing Solutions for Precise *Drosophila* Identification in Multi-Chamber Setups

Accurate identification of *Drosophila* individuals is essential for behavioral quantification given their small size. DrosoMating addresses this through three integrated video-processing features that enhance recognition fidelity in multi-chamber environments.

First, an effective recognition region function isolates individual wells as discrete analysis zones (Fig. 1A bottom), eliminating interference from adjacent chambers and background artifacts. Second, an image learning function adapts to phenotypic variability across strains. Operators designate three sample flies (Fig. 1B), enabling the algorithm to internalize strain-specific morphological signatures for improved detection. Third, a manual threshold adjustment function permits post-recognition optimization. Users visually validate automated identifications and dynamically refine sensitivity parameters (Fig. 1C), correcting false positives/negatives. Collectively, these features ensure robust tracking accuracy across diverse experimental setups and genetic backgrounds.

**Figure. 1.**
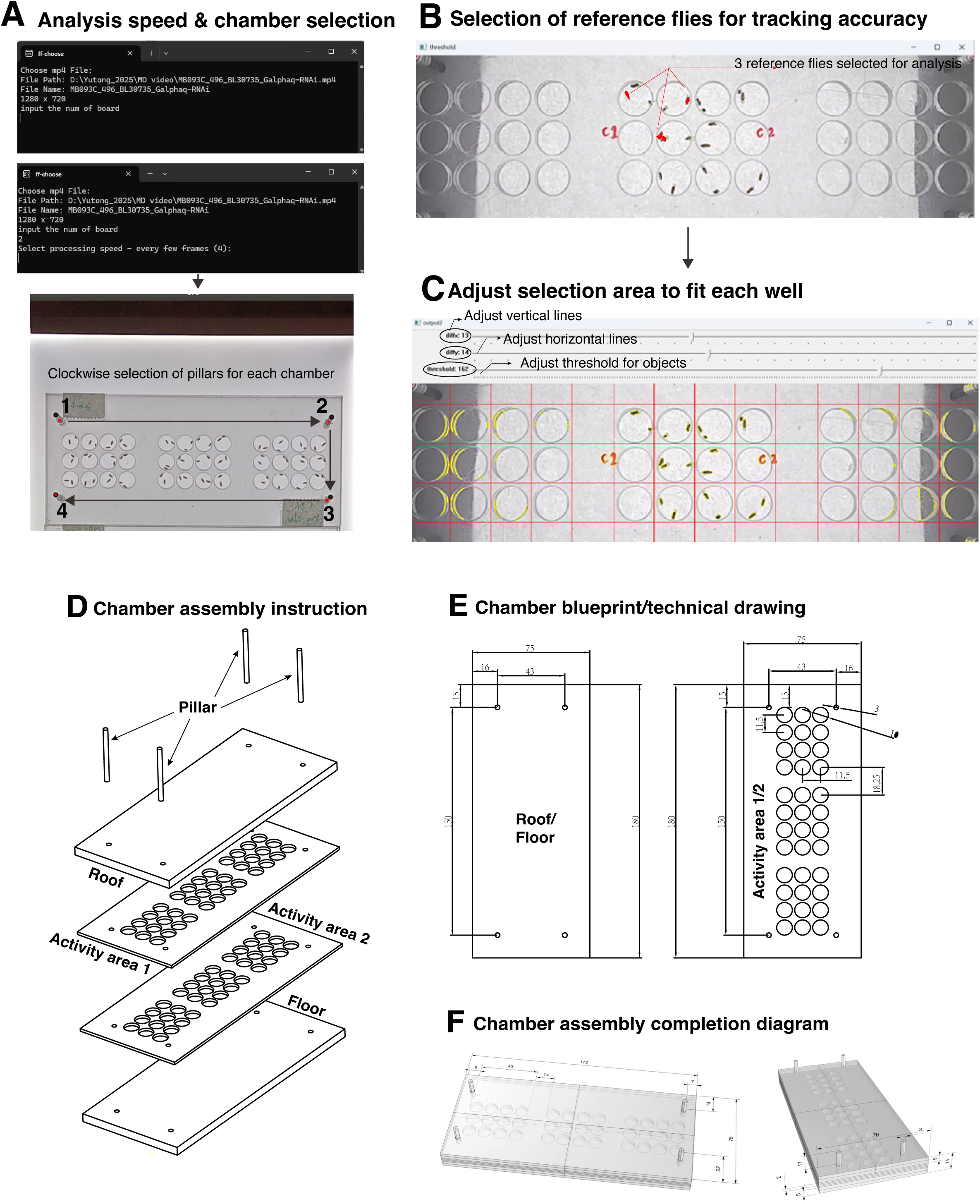
Usage of DrosoMating and CAD Parameters of the Chamber. **(A)** Typing input speed and input board numbers before selecting analyzing area (upper). Schematic representation of the region selection process (lower). Columns should be selected in a clockwise direction starting from the top left corner to define the chamber boundaries for perspective transformation (Fig. 1A, lower). Well numbering (1–36) follows a left-to-right, top-to-bottom sequence after perspective correction and is independent of pillar selection order. **(B)** Selection of reference flies for tracking accuracy. Any three reference flies were chosen for the analysis. **(C)** Use the X-axis, Y-axis, and threshold regulator to adjust the selection area to fit each well. Click “Enter” to run the program. Output file will be in the video folder: [Video Name]_output_[board numbers].csv (**D**) CAD exploded 3D view of the chamber instruction. (**E**) CAD parametric data of the chamber (dimensions in mm). (**F**) 3D diagram of the completed chamber

### Development and Validation of DrosoMating for Automated *Drosophila* Mating Behavior Analysis

To address the labor-intensive nature, high error rates, and limitations in quantifying complex behavioral metrics during manual observation of *Drosophila* mating, we developed the software DrosoMating. This tool automates analysis of mating videos recorded in a customized chamber. DrosoMating processes videos to generate a CSV file containing 6 key behavioral metrics: courtship duration, MD, CI, courtship start time, mating start time, and mating end time. While these metrics are individually straightforward, they can be combined to quantify diverse aspects of mating behavior metrics.

DrosoMating enables quantification of specific mating behaviors based on research needs. We focused on three core indices:

1. **Courtship Index (CI):** Indicates male courtship investment per unit time. Higher CI values (reflecting longer durations of behaviors like wing vibration or female chasing) suggest stronger male courtship motivation or higher female attractiveness (Greenspan and Ferveur, 2000b).
2. **Copulation Latency (CL):** Measures the time from courtship initiation to copulation start. Shorter CL indicates efficient male courtship or high female receptivity. Longer CL may reflect male courtship deficits (Eastwood and Burnet, 1977a), intense female rejection, or environmental stressors (Greenspan and Ferveur, 2000b; Singh and Singh, 2014).
3. **Mating Duration (MD):** Represents the copulation period length. Longer MD may enhance sperm competition but increases predation risk and energy costs, illustrating an evolutionary trade-off between reproduction and survival (Greenspan and Ferveur, 2000b).

Together, CI, CL, and MD provide a robust framework for characterizing *Drosophila* behavioral states during mating. Accurate quantification of these metrics is essential.

Because overlapping behaviors occur frequently during courtship and mating, DrosoMating handles short and prolonged overlaps differently. When flies overlap for ≤15 frames, centroid velocity continuity is used to assign identity; if overlap >15 frames, the region is flagged ‘occluded’ and excluded from CI calculation. Occluded frames are excluded from CI calculation but retained for mating-state detection via merged-contour analysis, ensuring continuous MD measurement through copulation.

We validated DrosoMating by comparing its output for CI, CL, and MD against values from repeated, corrected manual measurements. Differences between software-generated data and manual data were consistently within 10 seconds for all three metrics, with no statistically significant differences (Fig. 2A-C). Manual quantification of these metrics requires several hours of effort and is prone to substantial human error. In contrast, DrosoMating automates the analysis. After a few minutes of initial setup and configuration, the software generates precise behavioral metrics from mating videos (Fig. S3).

**Figure. 2.**
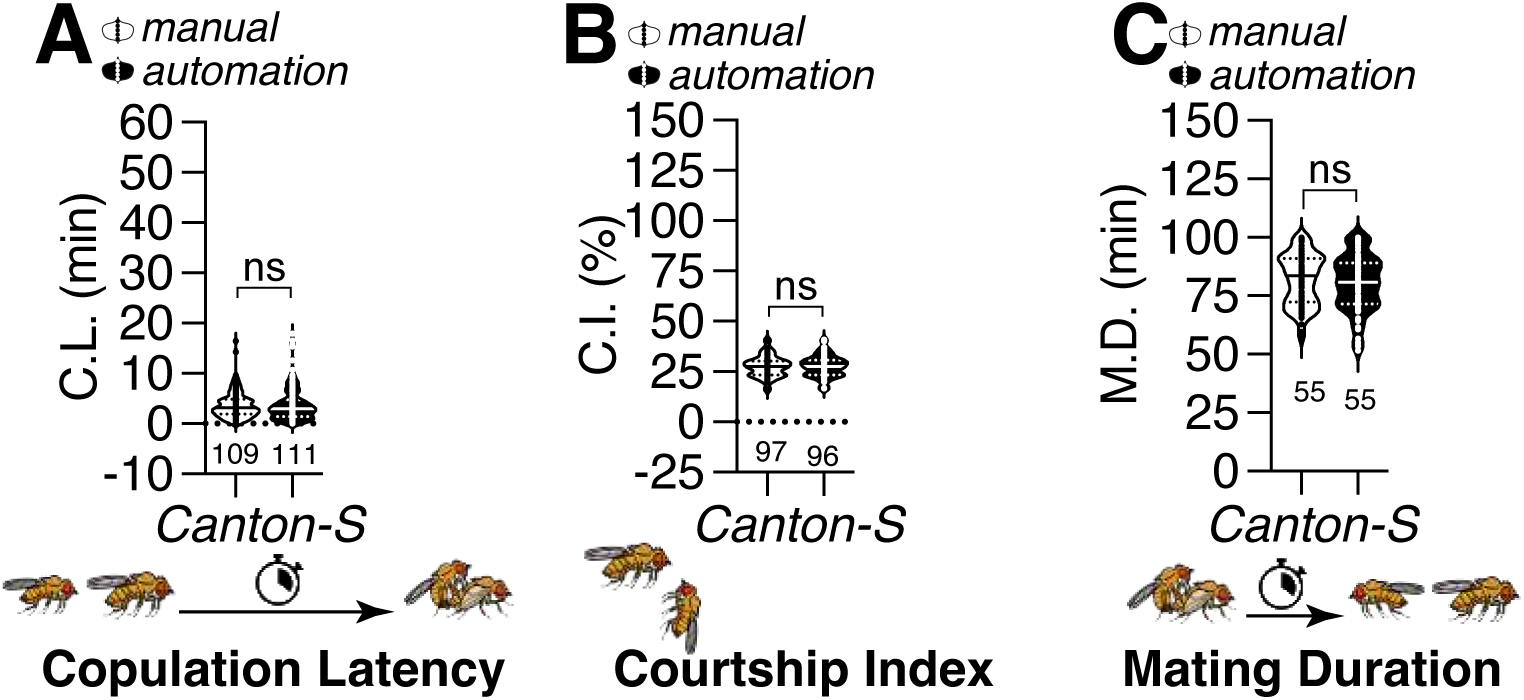
Comparison of the Accuracy of DrosoMating with Manual Scoring. **(A**) Copulation latency (CL) of Canton-S males was analyzed using DrosoMating and compared with manual recordings. For this figure, DBMs represent the ‘difference between means’ for the evaluation of estimation statistics. Asterisks represent significant differences, as revealed by the Student’s t test (*\* p<0.05, ** p<0.01, *** p<0.001*). Differences between methods were assessed using two-sided Student’s t-test (ns, not significant). See the Background for a detailed description of the CL assays used in this study. **(B)** Courtship index (CI) of Canton-S was analyzed by DrosoMating and manual recording. CI was measured by [courtship time/ (mating time-courtship time) *100%]. For detailed methods, see the BACKGROUND for a detailed description of the CI assays used in this study. **(C)** Mating Duration (MD) of Canton-S was analyzed by DrosoMating and manual recording. See the BACKGROUND for a detailed description of the MD assays used in this study.

### Validation of DrosoMating Across Diverse *Drosophila* Strains and Behavioral Paradigms

DrosoMating was adapted to quantify mating behavior metrics across diverse *Drosophila* strains and experimental conditions. To evaluate its universality and accuracy, mating interactions were analyzed between virgin females and males of four strains: Canton-S (wild-type), Oregon-R (wild-type), *w^1118^* (white eye mutant), and *y^1^* (yellow body color mutant). These lines have been extensively employed as standard inbred strains in research contexts for a prolonged period. Wild-type strains (Canton-S and Oregon-R) exhibited significantly higher CI, CL, and MD compared to *w^1118^* and *y^1^* mutants (Fig. 3A–C). This quantitative assessment revealed markedly reduced activity in mutant males across all three behavioral metrics, consistent with known phenotypic limitations: *w^1118^* males displayed restricted courtship due to visual impairment (Krstic et al., 2013), while *y^1^* males demonstrated suboptimal mating performance, aligning with prior reports of courtship deficits (Drapeau et al., 2006).

**Figure. 3.**
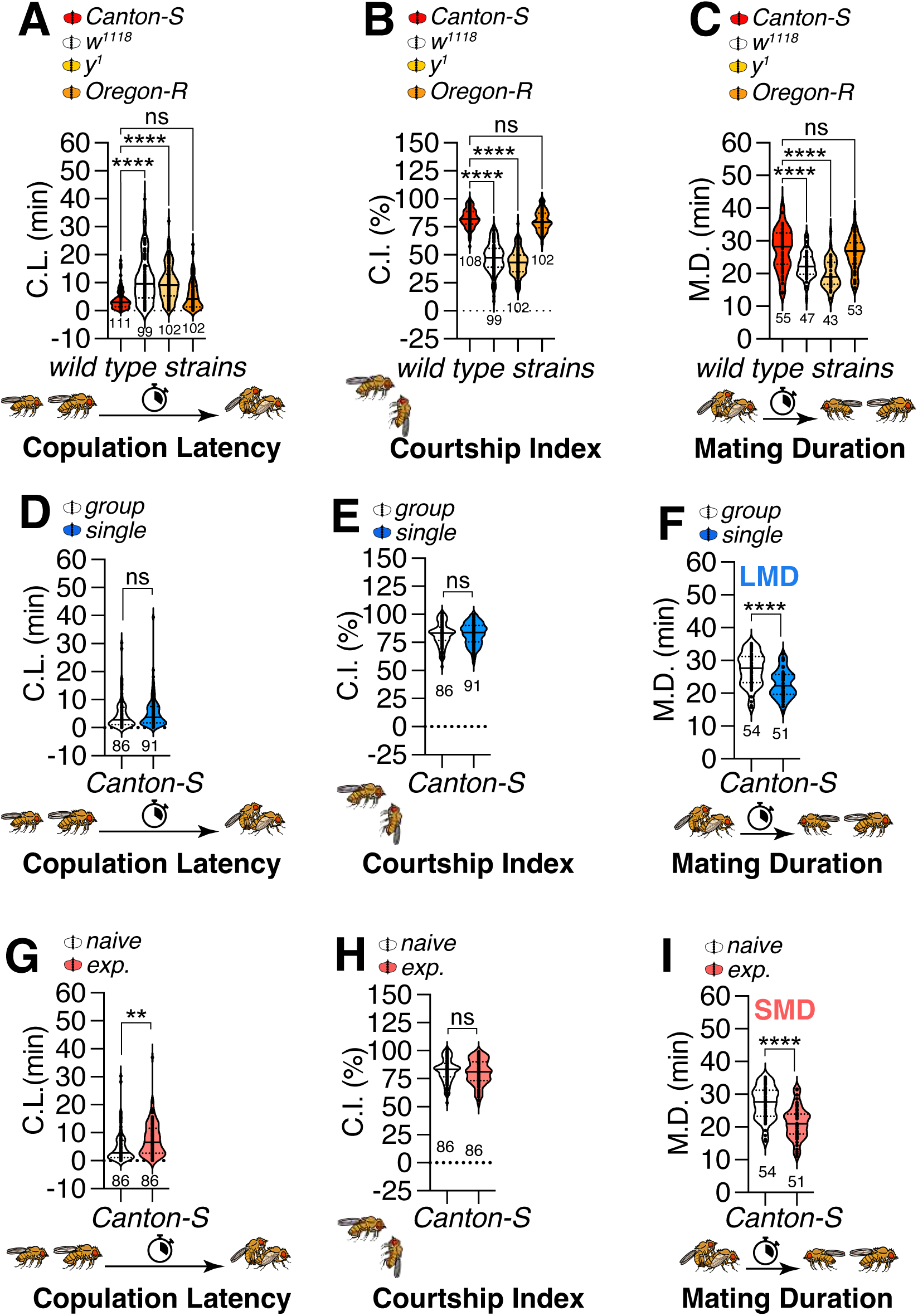
Behavioral assays of *Drosophila* males under different rearing conditions. **(A)** CL of Canton-S, *w^1118^*, *y^1^*, and Oregon-R males. **(B)** CI of Canton-S, *w^1118^*, *y^1^*, and Oregon-R males. **(C)** MD of Canton-S, *w^1118^*, *y^1^*, and Oregon-R males. For detailed methods, see the background for a detailed description of the mating assays used in this study. Violin plots show the distribution of copulation latency (CL, in minutes) for Canton-S, w^¹¹¹8^, y¹, and Oregon-R males. Sample sizes are indicated below each group. Statistical significance was assessed using one-way ANOVA followed by Tukey’s post hoc test for multiple comparisons for Fig. 3A-C. ****p < 0.0001; ns, not significant. **(D)** CL of group-housed and single-reared Canton-S males. For fig. 3D-I, DBMs represent the ‘difference between means’ for the evaluation of estimation statistics. Asterisks represent significant differences, as revealed by the Student’s t test (*\* p<0.05, ** p<0.01, *** p<0.001*). Differences between methods were assessed using two-sided Student’s t-test (ns, not significant). **(E)** CI of group-housed and single-reared Canton-S males. **(F)** LMD assays of Canton-S males. In the MD assays, white data points denote males that were group-reared (or sexually naïve), whereas blue data points signify males that were singly reared. The dot plots represent the MD of each male fly. The mean value and standard error are labeled within the dot plot. **(G)** CL of naive and sexually experienced Canton-S males. **(H)** CI of naive and sexually experienced Canton-S males. **(I)** SMD assays of Canton-S males. White data points represent sexually-naïve males and pink data points represent sexually-experienced ones.

Notably, reduced basal locomotor activity in *w*^¹¹¹8^ and *y¹* mutants has been well documented in previous studies, independent of courtship behavior (Drapeau et al., 2006; Krstic et al., 2013). Consistent with these reports, our tracking data show that single-housed w^1118^ and y^1^ males exhibit lower average velocity than Canton-S and Oregon-R controls (Fig. S4A). These general locomotor differences are insufficient to fully explain the observed courtship and mating timing phenotypes, indicating that additional courtship_related processes contribute to the observed behavioral differences.

The software further validated established behavioral paradigms—Longer Mating Duration (LMD) (Kim et al., 2013a, 2012a) and Shorter Mating Duration (SMD) (Lee et al., 2023a)—using Canton-S males subjected to distinct pre-experimental conditioning. DrosoMating accurately detected expected differences in CL, CI, and MD between LMD (group vs. single housing) and SMD (sexually naive vs. sexually experienced) conditions (Fig. 3D–I), confirming its precision in resolving complex, paradigm-specific mating dynamics.

### Quantification of Learning-Memory Dynamics and Neural Circuit Integration Using DrosoMating

DrosoMating facilitates the investigation of learning and memory consolidation in *Drosophila* through quantifiable behavioral metrics. The Learning Index (LI) was calculated as LI = (CI_naive − CI_trained) / CI_naive × 100%. (Fig. 4A), This metric provides a standardized measure of experience-dependent behavioral modulation. This transformation of abstract behavior into discrete parameters enables rigorous analysis of neural and behavioral plasticity.

**Figure. 4.**
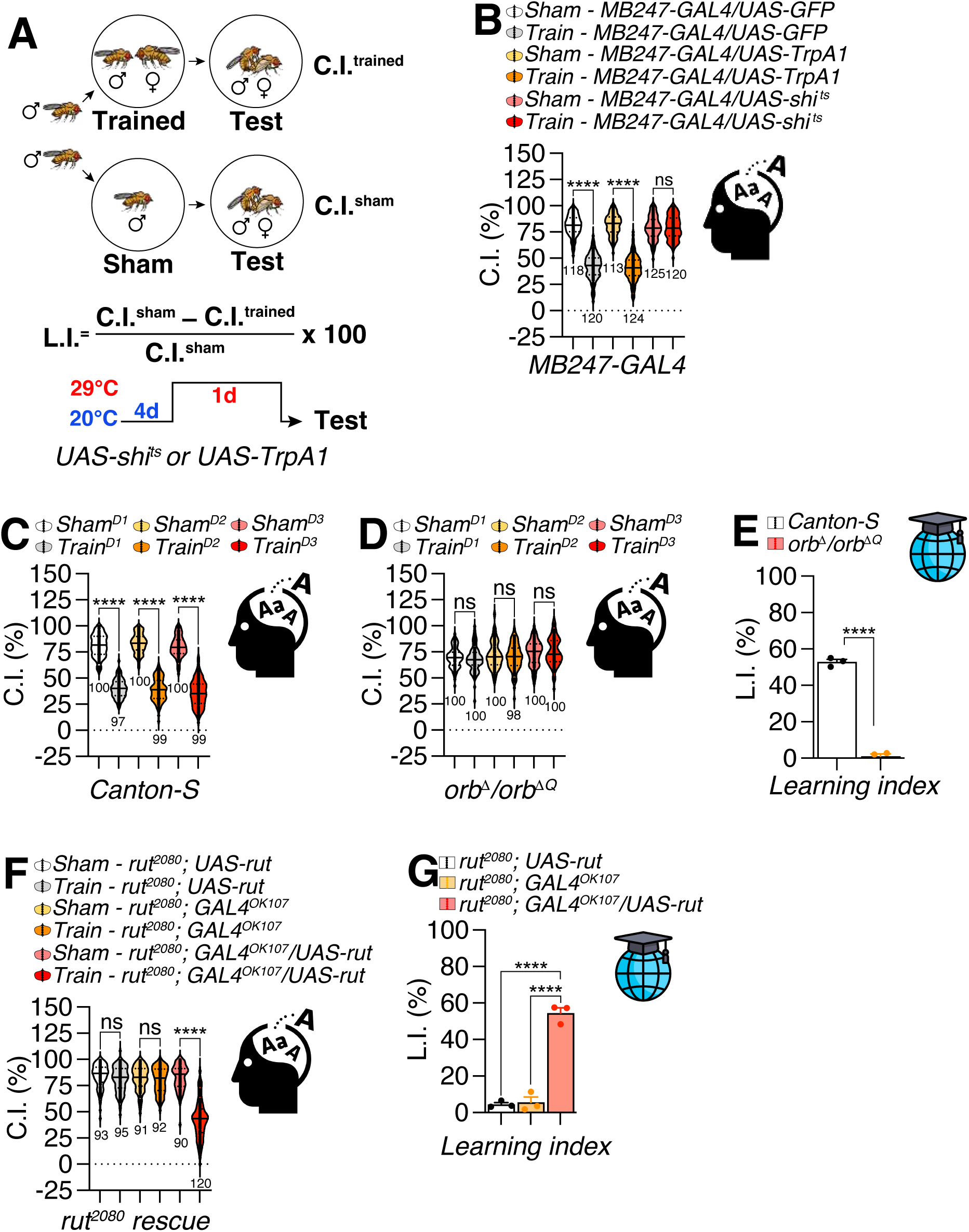
behavioral assays of learning in *Drosophila* which combined with temperature-dependent temporal activation/inhibition methods. **(A)** Learning Index (LI) calculation and experimental design for temperature-dependent activation/inhibition of neural circuits in adult *Drosophila*. This figure illustrates the experimental paradigm for calculating the LI using CI measurements from sham and trained group. The protocol involves rearing flies at 20°C for four days followed by a one-day exposure to 29°C to activate/inhibit temperature-dependent, genetically-encoded neuronal modulators (for example, *shi^ts^* as inhibitor or *TrpA1* as activator). **(B)** CI of training and sham training *MB247-GAL4/UAS-GFP*, *MB247-GAL4/UAS-TrpA1* and *MB247-GAL4/UAS-shi^ts^* flies. Sample sizes (n) are indicated below each group. Statistical significance was assessed using two-way ANOVA for Fig. 4 B,C,D and F, followed by Sidak’s post hoc test for pairwise comparisons between sham and trained groups within each genotype. ****p < 0.0001; ns, not significant. (**C**) CI of different training days (D1: 1 day, D2: 2 days, D3: 3 days) and sham trained Canton-S flies. (**D**) CI of different training days (D1: 1day, D2: 2 days, D3: 3 days) and sham training *orb*^Δ^*/orb*^Δ*Q*^ flies. (**E**) LI of Canton-S and *orb*^Δ^*/orb*^Δ*Q*^ males. For fig. 4E and G, DBMs represent the ‘difference between means’ for the evaluation of estimation statistics. Asterisks represent significant differences, as revealed by the Student’s t test (*\* p<0.05, ** p<0.01, *** p<0.001*). Differences between methods were assessed using two-sided Student’s t-test (ns, not significant). (**F**) CI of training and sham training *rut^2080^; UAS-rut, rut^2080^; GAL4^OK107^, rut^2080^; GAL4^OK107^/UAS-rut* flies. (**G**) LI of *rut^2080^; UAS-rut, rut^2080^; GAL4^OK107^, rut^2080^; GAL4^OK107^/UAS-rut* flies.

The software integrates seamlessly with neural manipulation techniques. For acute temporal control, thermogenetic tools (*UAS-shi^ts^* (Kitamoto, 2001), *UAS-TrpA1* (Kang et al., 2012)) permit precise activation or silencing of circuits during courtship (Fig. 4B and S1A). Targeted perturbation of mushroom body Kenyon cells, critical for memory processing, revealed immediate behavioral consequences (Fig. S1B–C), elucidating their role in behavioral adaptation.

Established courtship conditioning paradigms (Gil-Martí et al., 2023a; Griffith and Ejima, 2009a; Kamyshev et al., 1999; Reza et al., 2013; Wolf et al., 1998) are robustly replicated, validating DrosoMating’s fidelity in experience-driven behavior. (Fig. 4C). Mutant analyses revealed *orb2* (Fig. 4D–E) and *rut* (Fig. 4F-G) dependence of experience-induced CI reduction, consistent with known memory pathways.

Chronic neural interventions are enabled via *tub-GAL80^ts^*-mediated adult-specific silencing or knockdown (Fig. S1D–F). This approach reveals enduring impacts of sustained circuit modulation on courtship plasticity and memory consolidation. Collectively, DrosoMating bridges behavioral quantification and mechanistic inquiry, advancing exploration of learning-memory interplay through precise, scalable experimental frameworks.

### DrosoMating is more compatible with low-quality mating videos than conventional tracking-based pipelines

To evaluate whether conventional fly-tracking pipelines could be used as alternative tools for extracting mating-duration metrics, we tested Ctrax and FlyTracker on representative low-quality single-chamber mating videos recorded under our standard high-throughput conditions (Fig. 5). Ctrax is designed to estimate the position and orientation of multiple walking flies while maintaining individual identities over time (Branson et al., 2009) (https://ctrax.sourceforge.net/), whereas FlyTracker aims to track fly pose, including position, orientation, body size, wing and leg positions, and to generate trajectory- and feature-based outputs for downstream behavior analysis (Eyjolfsdottir et al., 2014). Because these programs were not readily compatible with our full high-throughput behavioral recording setup, we first cropped the original videos and tested single-chamber videos containing one male and one female fly.

**Figure 5.**
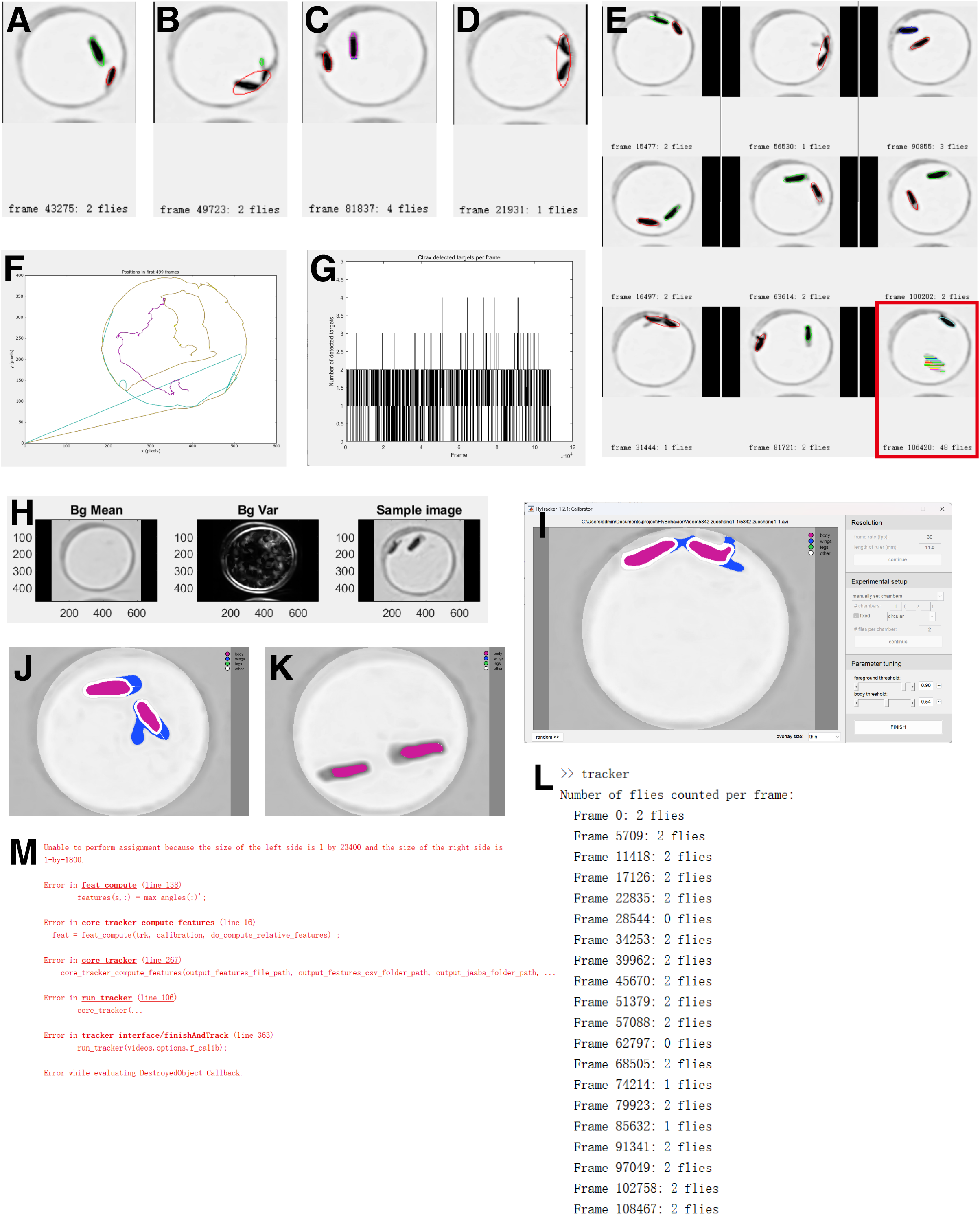
Limitations of conventional tracking-based tools for low-quality *Drosophila* mating videos. (A-D) Representative Ctrax detection frames from single-chamber mating videos, showing examples with different detected target numbers. (E) Additional randomly sampled Ctrax detection frames during tracking. (F) Representative Ctrax trajectory plot generated from the tracked video. (G) Frame-by-frame plot of the number of targets detected by Ctrax. (H) FlyTracker background model, background variance, and sample image used for detection. (I-K) Representative FlyTracker segmentation outputs under different threshold settings. (I) Segmentation output using the default threshold setting. (J) Example frame showing relatively successful fly segmentation under the default threshold setting. (K) Representative segmentation output after lowering the threshold to reduce tracking/feature-computation errors. (L) FlyTracker command-window output showing the number of flies detected in sampled frames. (M) Example FlyTracker error message during feature computation.

Using Ctrax, we observed that target detection was highly variable even within single-chamber videos. In representative frames, Ctrax could occasionally identify the two flies correctly (Fig. 5A). However, imperfect segmentation was frequently observed. In some frames, parts of the fly body were detected as additional targets, resulting in over-segmentation and an apparent increase in the number of detected flies (Fig. 5B,C). Conversely, when the male and female were close to each other or physically overlapped, the two animals were sometimes detected as a single target (Fig. 5D). These examples indicate that Ctrax detection was sensitive to the low contrast and overlapping fly bodies present in our mating videos. We then adjusted the Ctrax detection threshold to improve segmentation quality. Although threshold optimization improved detection in some frames, abnormal detections remained evident across randomly sampled frames (Fig. 5E). For example, some frames still showed incorrect target numbers, and a severe segmentation failure was observed in the lower-right example, where the detected objects did not correspond to two clearly separable flies. Thus, even after parameter optimization, Ctrax did not consistently maintain the expected two-target detection state in single-chamber mating videos.

The instability of Ctrax detection was also reflected in the trajectory output. In the first 500 frames, the generated trajectories were fragmented into multiple colored track segments rather than two continuous trajectories corresponding to the male and female (Fig. 5F). In addition, some trajectories extended outside the chamber boundary, indicating tracking errors and identity instability. Consistently, frame-by-frame quantification of detected target number showed that the detected object count did not remain stable at the expected value of two flies per chamber (Fig. 5G). Because Ctrax failed to maintain stable two-fly detection and continuous trajectories under these video conditions, its output could not be reliably used for downstream extraction of copulation latency or mating duration.

We next tested FlyTracker on the same type of cropped single-chamber mating videos. FlyTracker was developed to track multiple flies by estimating body position, orientation, size, wing and leg positions, and by maintaining fly identities across video frames; it also outputs per-frame features such as velocity, facing angle, and wing-angle-related measurements for downstream behavioral analysis(Eyjolfsdottir et al., 2014). In our videos, FlyTracker was able to generate a background model, indicating that the program could recognize the overall imaging field and chamber background (Fig. 5H). However, during calibration and segmentation, the default threshold setting produced inconsistent detection results (Fig. 5I). Although some frames were segmented relatively well under the default threshold (Fig. 5J), these successful examples were not representative of the overall tracking process, and the program frequently terminated with runtime errors during tracking or downstream feature computation. To improve detection stability, we lowered the segmentation threshold. Under this adjusted setting, FlyTracker produced more complete fly masks in representative frames (Fig. 5K), and the diagnostic output showed that two flies were detected in many sampled frames (Fig. 5L). Nevertheless, detection remained unstable in some sampled frames, including frames in which zero or one fly was detected despite the expected two flies per chamber (Fig. 5L). The full pipeline ultimately failed during feature computation, producing a runtime error before complete tracking and feature outputs could be generated (Fig. 5M). Therefore, even after threshold adjustment, FlyTracker could not provide a stable end-to-end workflow for extracting mating-duration metrics from these low-quality mating videos.

This failure mode is relevant because FlyTracker depends on stable segmentation, identity maintenance, and per-frame feature extraction. In our assay videos, low contrast, chamber-edge artifacts, and prolonged male–female overlap during copulation interfered with these requirements. As a result, FlyTracker could occasionally identify the flies in individual frames, but it did not reliably complete the full analysis pipeline required for downstream behavioral quantification. This limitation is especially important for workflows such as JAABA, which use manually labeled examples to train behavior classifiers but still depend on upstream tracking-derived features. Thus, for our low-quality high-throughput mating recordings, FlyTracker-based analysis was substantially less practical than DrosoMating, which directly outputs mating-related timing metrics without requiring continuous high-fidelity two-fly pose tracking.

## List of abbreviations

**abbreviations full term**

CL: Copulation Latency
MD: Mating Duration
CI: Courtship Index
LI: Learning Index
SMD: Shorter Mating Duration
LMD: Longer Mating Duration
CAD: Computer-Aided Design

## Methods and Materials

### Fly Stocks and Husbandry

*Drosophila* stocks were maintained on standard cornmeal-agar food at 25°C and 60-70% relative humidity, with a 12-hour light/dark cycle. When suppression of temperature-sensitive transgene activity was required (e.g., for *UAS-shi^ts^* or *UAS-TrpA1* experiments), flies were reared at 19-20°C throughout development. Because developmental duration at 19-20°C is approximately twice that at 25°C, experimental scheduling was adjusted to accommodate the delayed eclosion. Transgene activation was achieved by transferring flies to 29°C *(shi^ts^*) or 28°C (*TrpA1*) for 30 mins prior to experiments. To eliminate confounding variables in behavioral assays, control and experimental groups were always raised under identical environmental conditions, including temperature, humidity, and light cycle.

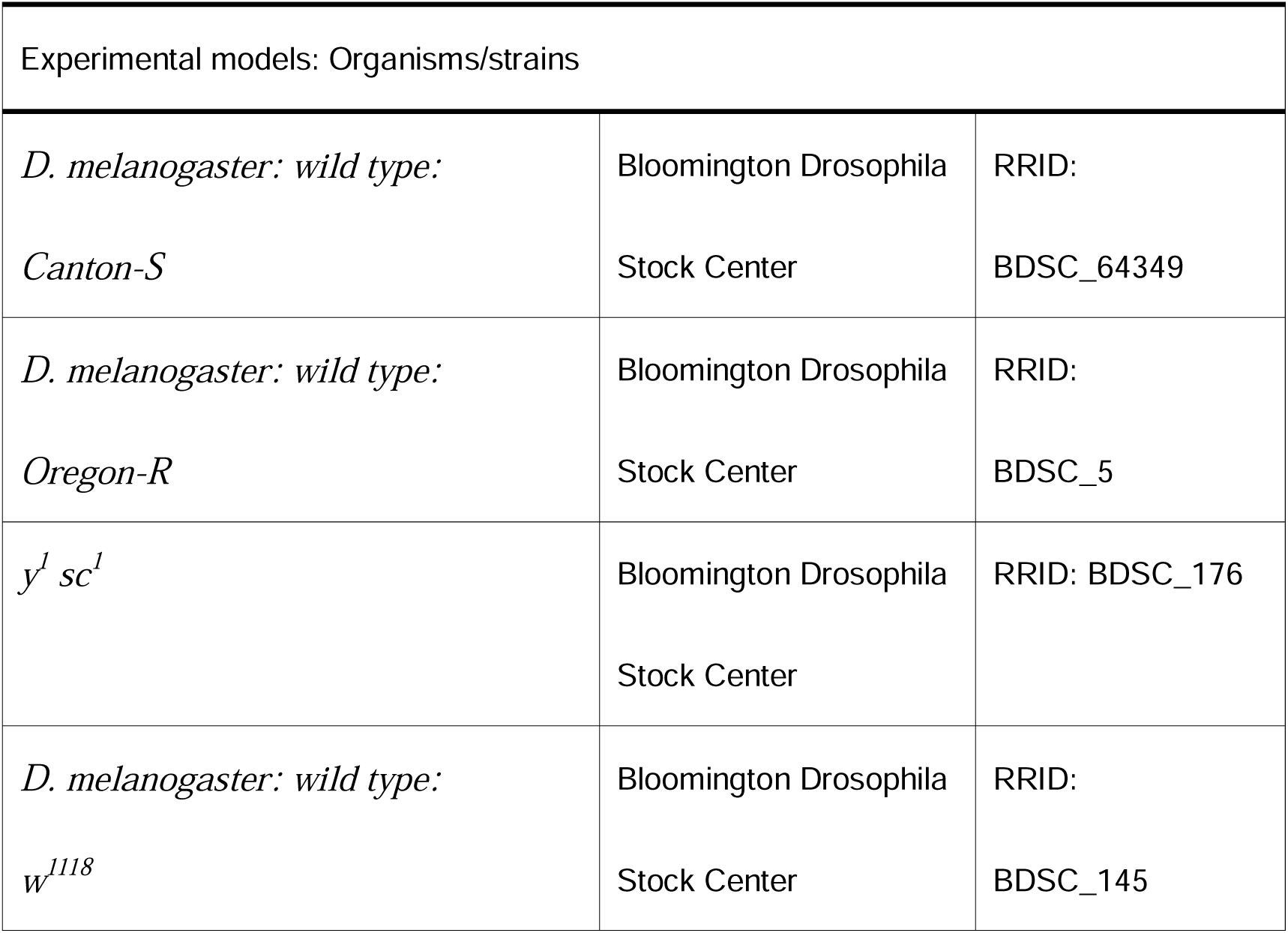
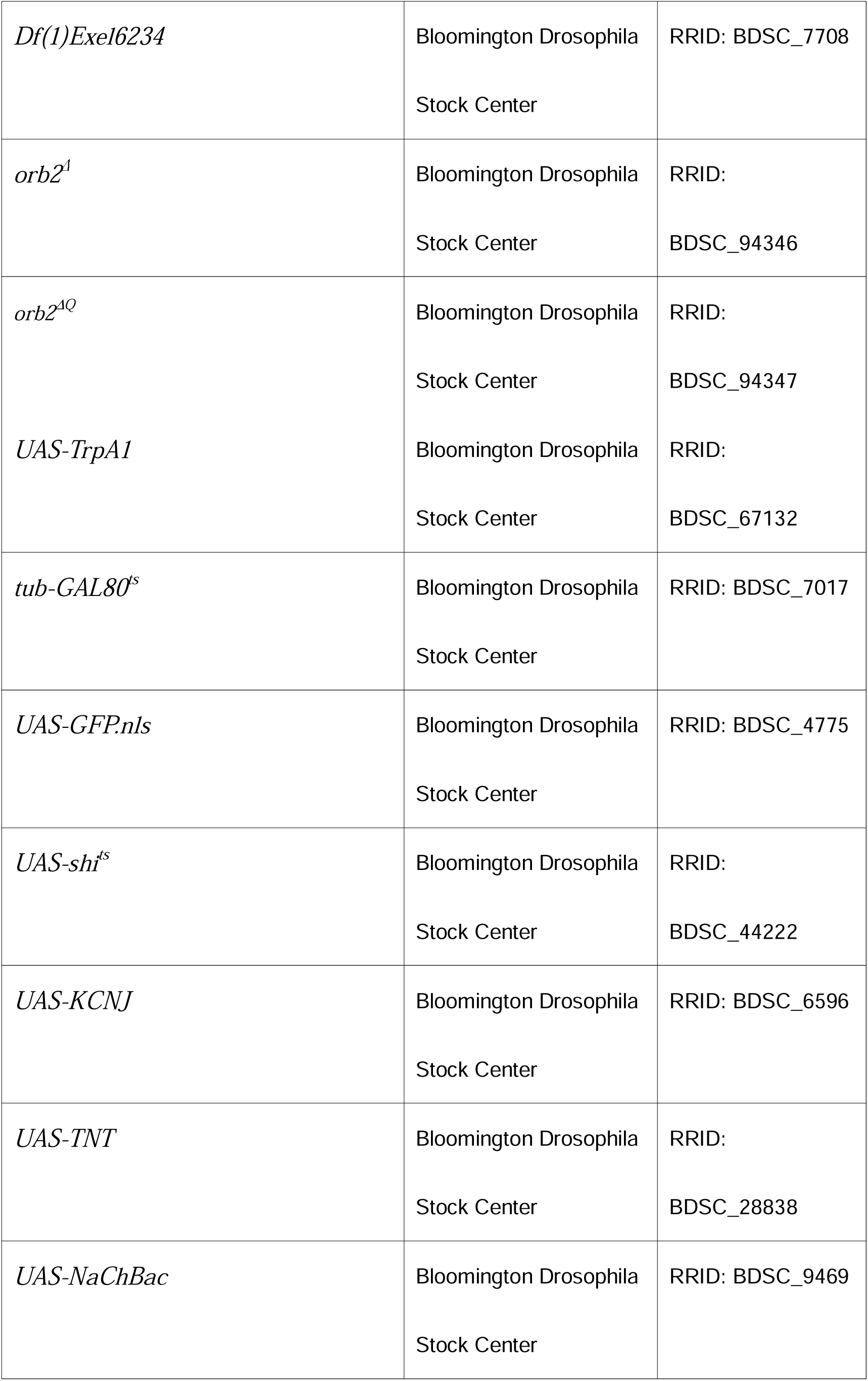
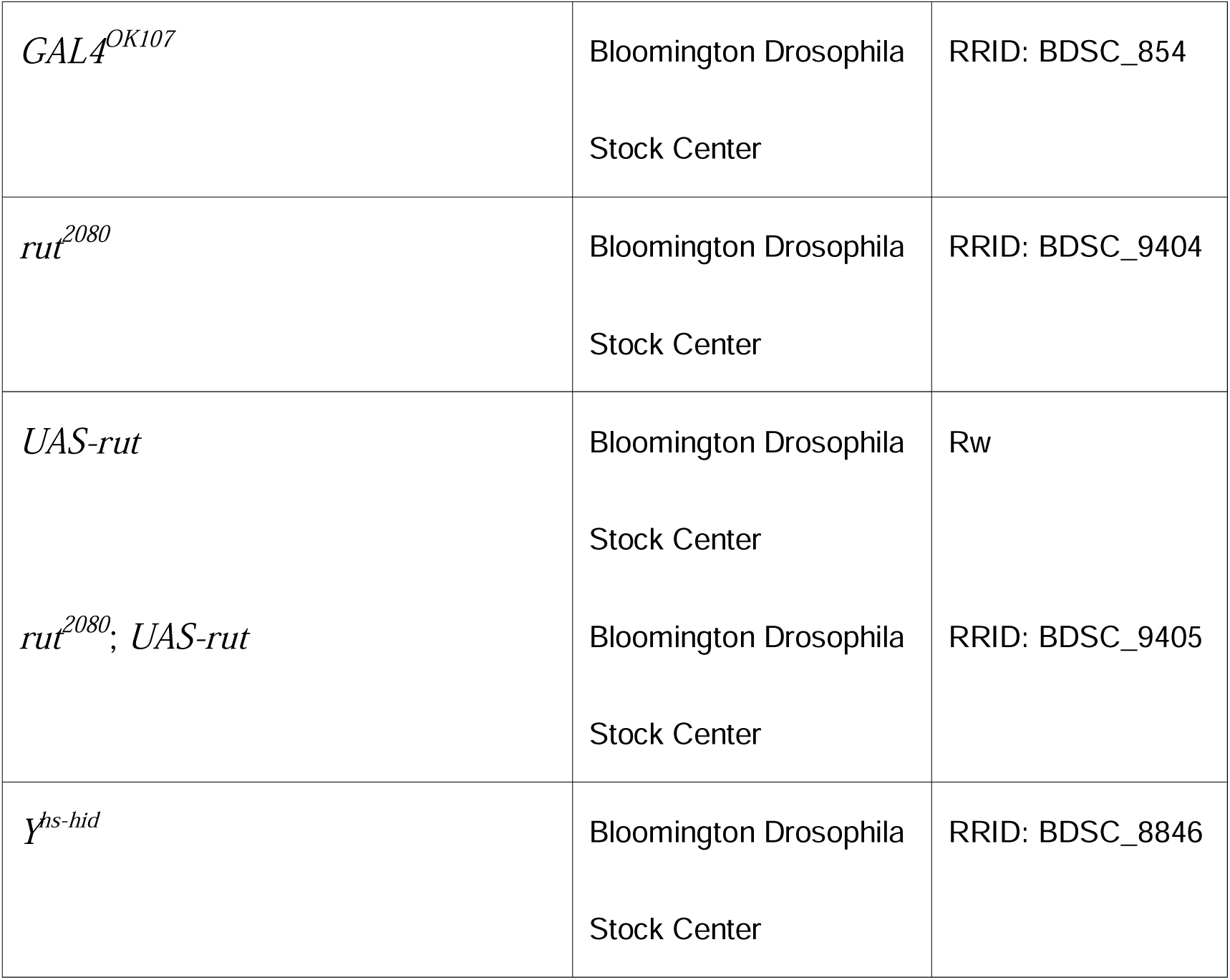

### Environmental Control System

The controlled-environment chamber is a dedicated fly room equipped to maintain standardized experimental conditions. It features a precision temperature control unit, which is a programmable apparatus for maintaining constant temperature with an accuracy of ±0.5°C. The chamber also includes automated humidity regulation through a humidifier or climate control system to stabilize relative humidity, typically within the range of 60–70%. Additionally, a programmable LED lighting system with full-spectrum arrays and an integrated timer is used to simulate circadian cycles, allowing for adjustable day/night duration and light intensity.

### Mating Behavior Measurement System Setup

1. **Behavioral observation arena**: The behavioral observation arena was constructed from transparent acrylic plates. Each chamber measured approximately 1 × 1 × 0.2 cm unless otherwise specified. (Fig. 1D-E).
2. **Sex-separation films**: Non-permeable acrylic partitions (0.5 mm thickness) to isolate male and female cohorts prior to trials. Customized apparatus components can be fabricated by precision cutting of OHP (overhead projector) film sheets.
3. **Uniform illumination system**: Programmable LED panel array (full-spectrum, 6500K color temperature) positioned to optimize contrast for automated tracking software.
4. **High-resolution imaging**: OPPO A72 5G main camera (FHD/1080p resolution, 30 fps) mounted on an adjustable stabilizer for consistent video capture.

### Mating Behavior Measurement Preparations

This protocol outlines the collection of male and virgin female *Drosophila melanogaster* and details the standardized mating behavior assay. To ensure experimental reproducibility, each cohort (genotype or condition) must include ≥100 individuals per sex, with final mating counts reaching at least 20 successful pairs (ideally 70-80) within a 2-hour observation window. The protocol spans developmental stages from egg to adult emergence, with adults requiring 5 days post-eclosion to attain sexual maturity. Flies are reared under controlled environmental conditions (25°C, 60-70% relative humidity, 12-hour light/dark cycle) to minimize physiological variability. To avoid confounding effects of anesthesia on mating behavior, CO_2_ exposure was strictly minimized during fly handling and sorting; brief, low-intensity exposure was reserved for initial virgin collection and before starting behavioral assays.

1. **Preparation of experimental male flies (15–20 days)**: Raise flies of the desired genotype (wild-type, mutants, or crosses such as GAL4/UAS combinations) in bottles at the specified temperature (standard: 25°C; alternative: 19°C or 29°C as required) under a 12:12 h light:dark cycle. Maintain 2–3 bottles per genotype, each containing 25 females and 15 males, to yield 25–30 flies per day. Transfer flies to fresh food every 2–3 days to prevent overcrowding.
2. **Post-eclosion handling**: Collect newly eclosed males (0–8 h post-eclosion) under brief CO□ anesthesia. Isolate individual males in vials and age them for 5 days at 25°C (or 9 days at 18°C) to ensure complete brain maturation. Clear existing adults from culture vials daily using CO□ to ensure synchronized collection of newly eclosed males. Group males into cohorts of 40 and rear in vials for 5–9 days to optimize sexual maturity while preserving mating efficiency.
3. **Large-scale virgin female collection for high-throughput genetic screening:** The *Df(1)Exel^6234^/Y^hs-hid^* (Bloomington Stock Center #7708 with #8846) or the *SPR^attP^*/*Y^hs-hid^* strain (Bloomington Stock Center #84576 with #8846) were used in this study. Both strains carry a null mutation in the *sex-peptide receptor (SPR)* gene (Yapici et al., 2008), which disrupts post-mating female refractoriness, thereby enhancing receptivity to remating and heat-inducible *hid* expression eliminating male progeny and balancer heterozygotes. Maintain cultures below 20°C prior to egg collection to prevent premature *hid*-induced lethality in *Y^hs-hid^* containing stocks. Higher temperatures (>20°C) may activate leaky *hs-hid* expression, reducing male lethality efficiency during subsequent heat shock.

Place adult *Df (1)Exel^6234^/Y^hs-hid^* flies in fresh bottles for 24 h to deposit eggs at 22°C. Heat shock incubate eggs at 37°C for 75 min to trigger *hid*-mediated apoptosis of males and balancer heterozygotes. We recommend using a water bath heated to 37°C for efficient heat shock instead of simply placing the samples in a 37°C incubator. Collect surviving female progeny and rear in groups of 40 for 5–8 days under standard conditions at 25°C to produce age-matched, virgin females. Raising at 29°C can speed up the process of obtaining virgin females. Following heat shock treatment, inspect all eclosed adults to confirm complete absence of male progeny. The *Df(1)Exel^6234^/Y^hs-hid^* system should yield 100% female populations when properly executed.

#### 4. Chamber specifications

– **Structure:** Reinforced with four stainless steel columns (3 mm diameter) to ensure stability (Fig. 1D-E and 6A).
– **Material**: Four transparent acrylic plates (180 × 75 mm) with alternating thicknesses (1 cm and 0.3 cm), assembled in a “thick-thin-thin-thick” sequence (Fig. 1F and 6C).
– **Layout**: 36 individual chambers arranged in 3×4 clusters (12 chambers per cluster).
– **Chamber spacing**: 1 mm between individual chambers; 2 cm between clusters. Positioned 3 cm from the chamber’s long edge and 1 cm from the short edge (Fig. 1E-F).

Chamber dimensions are not fixed. By simply adjusting the magnification of the recording lens, users can re-scale the field of view and then refine the x (horizontal) and y (vertical) grid values in the DrosoMating interface to enclose the entire fly activity area within the corresponding wells. After this one-step calibration, the pipeline quantifies mating metrics (CL, CI, MD, etc.) with identical accuracy across different chamber sizes.

**Figure. 6.**
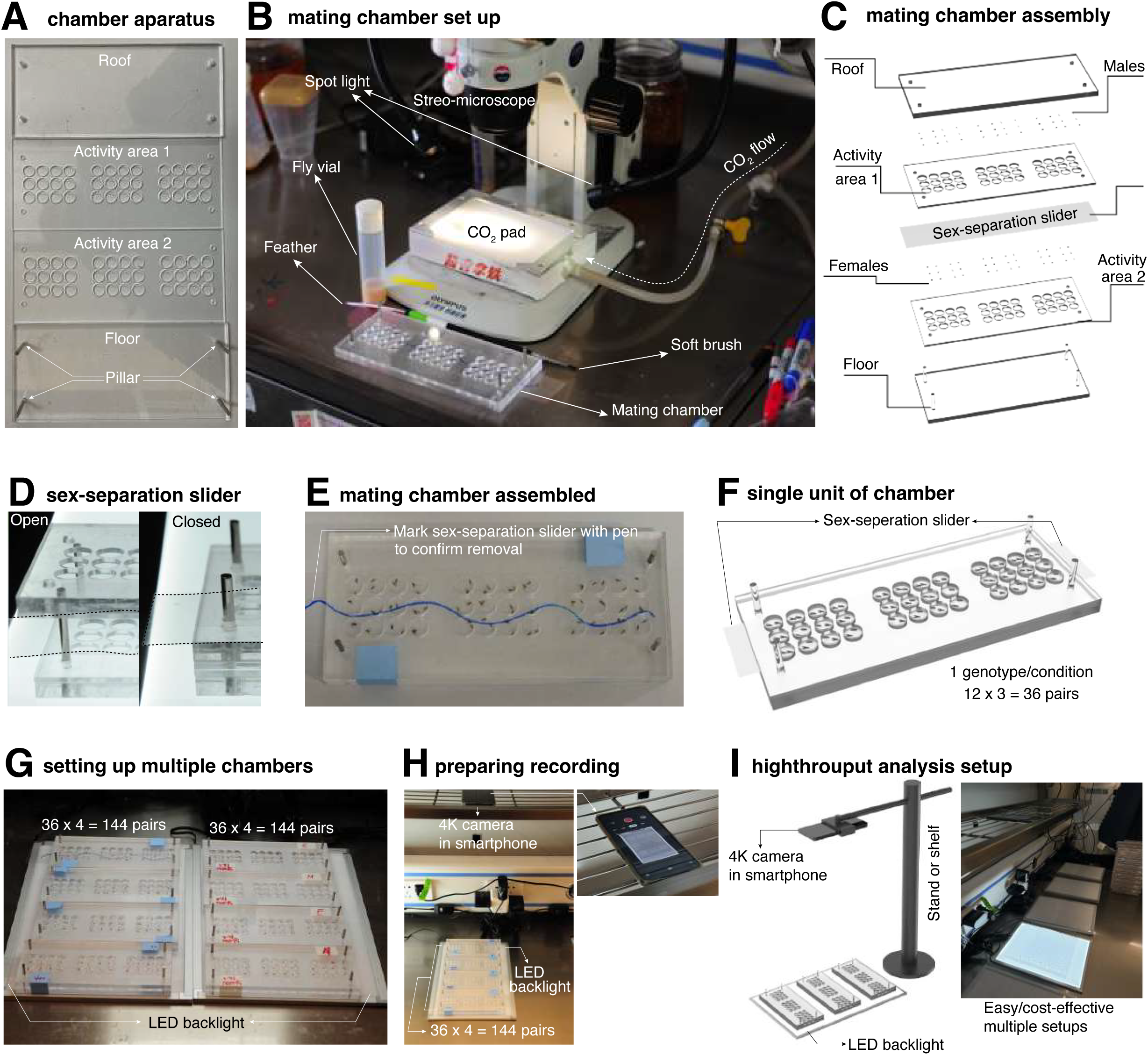
Setup of mating assay chamber apparatus and architecture of workstation. (**A**) Disassembly of the chamber apparatus. Each chamber comprises a roof, two activity areas, and a floor integrated with four pillars. (**B**) Workstation for assembling mating assay chambers. Essential required equipments for chamber assembly are shown: a stereomicroscope with an integrated light source, carbon dioxide anesthesia apparatus, a soft brush with feather on the opposite end, chamber apparatus, and fly containing vials. (**C**) 3D schematic of the mating chamber assembly. Assembly from bottom to top as shown. Briefly, first assemble the floor with one activity area 2 for placing females, then cover the females and separate activity area 1 from activity area 2 using the sex-separation slider. Next, position males and secure the chambers with paper tape. (**D**) The sex-separation slider is positioned between activity area 1 and activity area 2. (**E**) The mating chamber assembled with paper tape; the sex-separation slider should be marked with a pen to prevent forgetting to remove it. (**F**) 3D schematic of single chamber unit, can hold up to 36 pairs of fruit flies simultaneously. (**G**) Multiple chambers are set up with four chambers vertically aligned on an LED backlight, allowing the accommodation of 144 pairs of fruit flies across all chambers. (**H**) Video recording setup. Position a 1080p resolution camera or a smartphone with matching video capabilities above the chambers, turn on the LED backlight, and remove sex-separation slider before recording. (**I**) Setup for recording mating behavior using a camera mounted on a stand, positioned vertically above chambers arranged on a size-matched LED backlight to ensure uniform illumination. The camera is aligned to capture the entire chamber array, enabling simultaneous recording of mating interactions under controlled lighting conditions.

#### 5. Software implementation

Automated analysis is performed using the DrosoMating pipeline (<u>GitHub</u> https://github.com/hcls-kimlab/DrosoMating) (“https://github.com/hcls-kimlab/DrosoMating,” n.d.), the raw code is under win-ff module. which extracts courtship and mating metrics from video recordings. (Linux version: https://github.com/liuxiao916/Fruit_Flies)

### Mating Behavior Assay Step-by-step Procedures

1. **Fly preparation**: On day 5 post-eclosion, select healthy males (of the desired genotype) and virgin females. Lightly anesthetize flies using ice or CO□ pad (avoid prolonged exposure to prevent stress) (Fig. 6B). To minimize behavioral perturbations, flies should only be anesthetized at this step. For brief immobilization (e.g., sorting or transferring individuals), use CO□ anesthesia (≤30 s exposure). To avoid bias, all experiments should be done blindly. Label fly tubes with random numbers instead of genotypes and keep a separate list matching numbers to genotypes. Only check which numbers correspond to which genotypes after finishing all data analysis.
2. **Chamber assembly and fly loading**: Using a soft brush, gently place a single female into a chamber hole in Floor (Activity Area 2) (Fig. 6B). Cover the chamber with a transparent film to prevent physical contact while allowing visual/chemical interactions. Align Activity Area 1 on top of the film (Fig. 6C). Introduce a single male into the opposite side of the film using the transparent film as a slider to minimize CO□ exposure (Fig. 6D). Limit male exposure to CO□ during this step to avoid behavioral artifacts. Seal the chamber securely with Roof using disposable paper tape (Fig. 6E).
3. **Recovery and Mating Initiation**: Transfer the assembled chamber to a 25°C incubator (humidity controlled at 60–70% RH) for 90 min to ensure flies recover fully from handling stress. After recovery, gently remove the transparent film to initiate physical mating (Fig. 6E and F).
4. **Video Recording Setup**: Position an adjustable LED panel beneath the chamber to ensure uniform illumination (Fig. 6G). Record mating behavior at 1080p resolution for 1.5–2 h (adjust duration as needed) (Fig. 6H). A minimal setup for capturing fruit fly mating behaviors can be rapidly established without extensive external facility requirements. The essential components include a camera stand, an LED light panel, and a camera capable of recording at a resolution and frame rate of 1080p (30 fps). (Fig. 6I).

### Analysis of *Drosophila* Mating Video Data Analysis and Statistics

#### The use of the Windows version of DrosoMating

1. Prepare computer equipped either Windows (or Linux) PC with DrosoMating (ff-chose.exe) installed. The original code instructions for use are provided under the win-ff module.
2. Transfer video files into the computer to be analyzed. Launch DrosoMating and open the video file.
3. Enter the number of boards and velocity parameter. After entering the information, click “Enter” to proceed to the next step. (Fig. 1A)
4. Select the four steel marker pillars in clockwise order (Fig. 1A). Adjust the selection sequence based on the video orientation. For each plate, a preview image will appear. Click on three distinct flies to enhance tracking accuracy (Fig. 1B).
5. After selecting all regions, return to the DrosoMating home interface. Adjust x (horizontal) and y (vertical) values to align grids with chamber holes. Fine-tune fly detection by modifying the s value (sensitivity threshold) until all flies are correctly identified. Use the grid to segment individual flies (as shown in Fig. 1C). Note on the s value: The parameter s represents the grayscale intensity threshold (range: 0–255 for 8-bit images) used for binary segmentation of flies from the background. It is automatically calculated as the maximum grayscale value at the three reference fly positions plus an offset of 28, and can be manually adjusted to accommodate varying illumination conditions.
6. Run the analysis (Save the processed video but not recommended for routine use).
7. Export raw data as a CSV or Excel file. Open it in spreadsheet software for further processing.

**The usage instructions for the Linux version of DrosoMating are presented in Fig. S2**

### Statistical Analysis

To ensure robust statistical analysis, each experimental group included at least 100 male flies (naïve, sexually experienced, or singly reared). Internal controls were incorporated into every experiment as recommended by Bretman et al. (2011) (Bretman et al., 2011). Normality of the mating duration data was confirmed using the Kolmogorov-Smirnov test (*p*>0.05). For group comparisons, two-sided Student’s *t*-tests were applied to calculate significance levels (\*\*\*\**p*<0.0001, \*\*\**p*<0.001, \*\**p*<0.01, * *p*<0.05), comparisons among three or more groups were performed using one-way ANOVA with Tukey’s HSD post-hoc tests. Two-factor experimental comparisons were performed using two-way ANOVA followed by Sidak’s post-hoc test. Estimation statistics (Claridge-Chang and Assam, 2016) were additionally used to visualize effect sizes, mean differences, and precision, avoiding reliance solely on null hypothesis testing. All analyses, including data plotting, were performed using GraphPad Prism software.

## Discussion

### Advancing Courtship Behavior Analysis in *Drosophila*: From Manual Tracking to computer-vision classifier

The study of male *Drosophila* mating behavior represents a cornerstone in neurogenetics, providing critical insights into the neural and molecular mechanisms underlying innate social behaviors (Dauwalder, 2008; Dukas, 2020). Historically, the CI—quantifying the proportion of time spent in courtship activities—has served as a fundamental metric for evaluating mating activity (Greenspan and Ferveur, 2000a; Griffith and Ejima, 2009b; Pavlou and Goodwin, 2013; Sokolowski, 2001; Yamamoto and Koganezawa, 2013). However, traditional methodologies relied on labor-intensive manual observation, limiting their capacity to capture dynamic behavioral repertoires such as orientation, wing extension (“courtship song”), and copulation sequences (Dauwalder, 2008; Dukas, 2020). While machine vision systems like Dankert et al.’s automated platform advanced quantification of social interactions, their primary application skewed toward male-male aggression rather than nuanced courtship dynamics (Dankert et al., 2009b). Concurrently, research into genetic and environmental modulators—including *fruitless* (*fru*) and *doublesex* (*dsx*) gene pathways, cuticular hydrocarbons, and sensory cues—highlighted the need for scalable, high-resolution tools to dissect multimodal influences on behavior.

### Technical Innovation and Validation

DrosoMating addresses these gaps through an integrated framework combining modular environmental chambers, automated video tracking, and machine-based annotation. This system minimizes manual intervention while capturing temporally precise metrics such as CL, CI, and MD. Validation against ground-truth manual scoring demonstrated exceptional accuracy (98–99% agreement; error rates <0.05%), underscoring reliability for high-throughput screens. Crucially, the platform detected subtle behavioral variations across genotypes—e.g., diminished courtship vigor in w^1118^ and y^1^ mutants—and environmental manipulations such as group-rearing or prior sexual experience, which respectively shortened mating duration by ∼8–12 %. This sensitivity enables investigations into adaptive trade-offs, learning phenotypes, and circuit-level mechanisms. For instance, DrosoMating replicated classic deficits in aversive learning following mushroom body ablation and courtship impairments in *rutabaga* learning mutants, corroborating its utility for behavioral neuroscience (Levin et al., 1992).

### Limitations and Future Directions

Despite its advantages, DrosoMating currently lacks granularity for posture-specific interactions (e.g., “wing threat” during aggression or “circling” during courtship) and multisensory integration. Future iterations could incorporate 3D pose estimation algorithms (e.g., DeepLabCut) or integrate JAABA-based classifiers to expand action repertoires (Kabra et al., 2013). Although DrosoMating does not currently export the full set of per-frame kinematic features required for direct JAABA classifier training, its modular video-processing framework could be extended in future versions to generate JAABA-compatible trajectory or feature outputs.

Cross-species testing was not performed in this study. Although DrosoMating’s state-detection approach is morphology-agnostic and therefore theoretically applicable across *Drosophila* species, we have revised the text to avoid claiming demonstrated cross-species performance. Formal validation across diverse species remains a promising future direction.

### Comparative Evaluation with Existing Courtship Analysis Tools

A major advantage of DrosoMating is its compatibility with low-quality, high-throughput mating videos. Conventional tracking-based tools such as Ctrax and FlyTracker are powerful for trajectory- and pose-based behavioral analysis, but they generally require stable object segmentation, identity maintenance, and reliable feature extraction across frames. Ctrax was designed to estimate the positions and orientations of multiple walking flies while maintaining their identities, whereas FlyTracker aims to track detailed fly pose and generate per-frame behavioral features. Under our recording conditions, these requirements were difficult to satisfy because the videos were low contrast and the male and female frequently overlapped during copulation.

In our tests, both tools showed limited compatibility with these videos. Even after cropping to single-chamber videos and adjusting detection parameters, Ctrax produced unstable target numbers, fragmented trajectories, and tracking errors. FlyTracker could generate a background model and occasionally segment flies successfully, but detection remained unstable and the full pipeline failed during feature computation. These issues are particularly relevant for copulation latency and mating duration analysis: during copulation, the male and female remain physically coupled for a long period, which makes identity-based tracking difficult. For these timing metrics, it is more important to robustly detect the onset and offset of the mating state than to reconstruct detailed individual trajectories. DrosoMating was purpose-built to address these specific challenges. It operates reliably on lower-quality video streams, requires no complex pre-processing or manual ROI definition, and is optimized for high-throughput multi-chamber analysis. While it does not offer the same level of pose or kinematic detail as other tools, it provides a unique solution for laboratories seeking a simple, fast, and robust pipeline to quantify core reproductive timing metrics—copulation latency (CL), courtship index (CI), and mating duration (MD)—without the overhead of more complex systems (Table.1).

**Table 1.** Comparison of DrosoMating with tracking-based tools under low-quality mating-video conditions. The comparison was performed using representative low-quality, high-throughput mating videos recorded under standard experimental conditions. DrosoMating is compatible with low-contrast video streams, supports native multi-chamber analysis (up to 4 boards, 144 wells total), and robustly handles copulation overlap via state-detection without requiring individual identity maintenance. It directly outputs copulation latency (CL), courtship index (CI), and mating duration (MD) as automated CSV files with high (98–99%) agreement relative to manual scoring. By contrast, Under the tested low-quality mating-video conditions, Ctrax and FlyTracker did not provide stable end-to-end outputs suitable for reliable extraction of mating-timing metrics. DrosoMating was designed to extract mating-related timing metrics from high-throughput videos without requiring continuous two-fly identity tracking. Ctrax and FlyTracker were tested on cropped single-chamber videos from the same recording setup. Their limitations described here refer specifically to these low-quality mating videos and should not be interpreted as general limitations of the tools.

|  | **DrosoMating** | **Ctrax** | **FlyTracker** |
| --- | --- | --- | --- |
| Performance on low-quality mating videos | Compatible | Unstable under tested conditions<br>(require high contrast) | Unstable under tested conditions<br>(require high contrast) |
| Multi-chamber analysis | Up to 4 chambers (36×4 wells) | Requires cropping or additional preprocessing | Requires cropping or additional preprocessing |
| Copulation overlap handling | Robust (state detection, no identity needed) | Unstable during overlap<br>(over-/under-segmentation) | Unstable during overlap<br>(Runtime error under tested videos) |
| Direct output of CL/CI/MD | Yes; automated CSV | No (unstable trajectories) | No (runtime error) |
| Accuracy vs. manual scoring | High agreement with manual scoring | Not quantified because no stable CL/MD output was obtained | Not quantified because no stable CL/MD output was obtained |

Although we directly tested only Ctrax and FlyTracker, this limitation may also affect workflows that depend on upstream tracking-derived features. For example, JAABA uses tracking-derived features to train behavior classifiers, and DANCE, a recent *Drosophila* aggression and courtship pipeline, uses JAABA-based classifiers and lists FlyTracker and JAABA as required software. Thus, our conclusion is not that these tools are generally unsuitable for *Drosophila* behavior analysis, but that DrosoMating provides a more practical workflow for low-quality, high-throughput videos focused specifically on mating timing.

While DrosoMating currently prioritizes mating timing metrics over discrete behavioral classification, its modular architecture provides a foundation for future integration with behavior classifiers such as JAABA. Laboratories requiring granular behavioral elements—such as wing extension or circling—would benefit from an extended pipeline that exports per-frame kinematic features for downstream classifier training. Validating this integration represents a promising future direction to broaden the tool’s utility beyond core reproductive timing assays.

## Data and code availability

Strains are available upon request. The authors affirm that all data necessary for confirming the conclusions of the article are present within the article, figures, and tables. Our software is freely available at <u>GitHub</u> https://github.com/hcls-kimlab/DrosoMating. (Linux version: https://github.com/liuxiao916/Fruit_Flies)

## Resource availability

### Lead contact

Further information and requests for resources and reagents should be directed to and will be fulfilled by the lead contact, Woo Jae Kim

### Materials availability

This study did not generate new, unique reagents.

## Acknowledgements

The authors thank Kim lab members for their kind advice on this manuscript. This work was supported by Startup funds from The HIT Center for Life Science (HCLS) to Woo Jae Kim.

## Author information

### Authors’ contributions

XL, YS, and DS conceived and designed the study. WK designed the chamber. XL and FJ completed the development of the software underlying code. YS, HM conducted the subsequent software development and testing. YX fabricated all the hardware devices. YS performed the DrosoMating analysis. WK, YS, HM, and DS wrote the manuscript. WK, YS, HM and DS created the figures and tutorial video. YS and HM revised the manuscript. All authors read, reviewed, and approved the final manuscript.

---

**Figure. S1.**
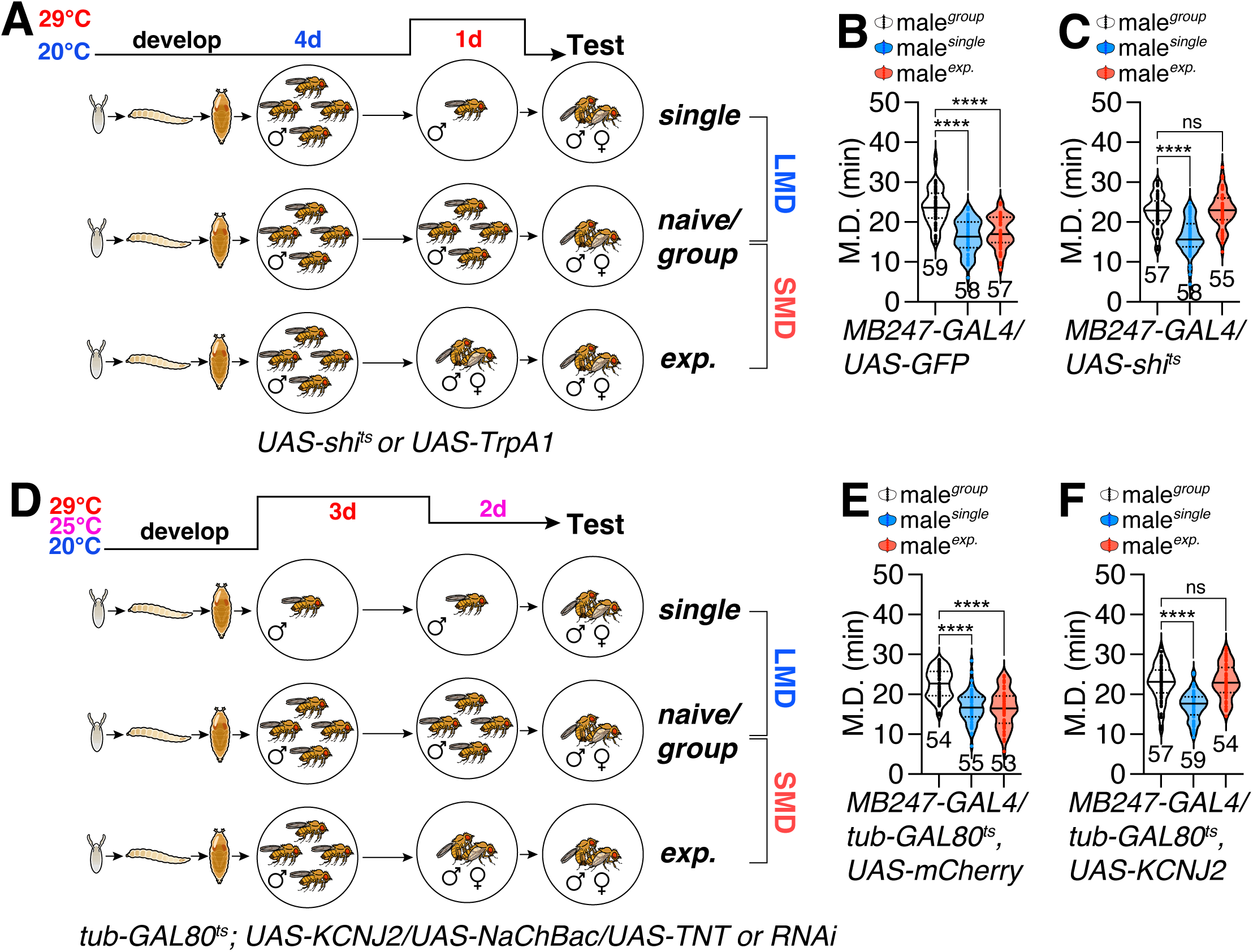
Temperature-dependent neural manipulation during LMD and SMD assays in adult *Drosophila*. **(A)** Schematic representation of LMD and SMD assays timeline when flies were crossed with heat-sensitive *Drosophila* cation channel *TrpA1* and *shi^ts^*. **(B)** MD assays of flies expressing the *MB247-GAL4* driver together with *UAS-GFP*. Sample sizes are indicated below each group. Statistical significance was assessed using one-way ANOVA followed by Tukey’s post hoc test for multiple comparisons for Fig. S1B-C,E-F. ****p < 0.0001; ns, not significant. **(C)** MD assays of flies expressing the *MB247-GAL4* driver together with *UAS-shi^ts^*. **(D)** Schematic representation of LMD and SMD assays timeline when *tub-GAL80^ts^; UAS-KCNJ2/UAS-NaChBac/UAS-TNT* or *RNAi* flies are crossed with specific GAL4 driver. **(E)** MD assays of flies expressing the *MB247-GAL4* driver together with *tub-GAL80^ts^*, *UAS-mCherry*. **(F)** MD assays of flies expressing the *MB247-GAL4* driver together with *tub-GAL80^ts^*, *UAS-KCNJ2*.

**Figure. S2.**
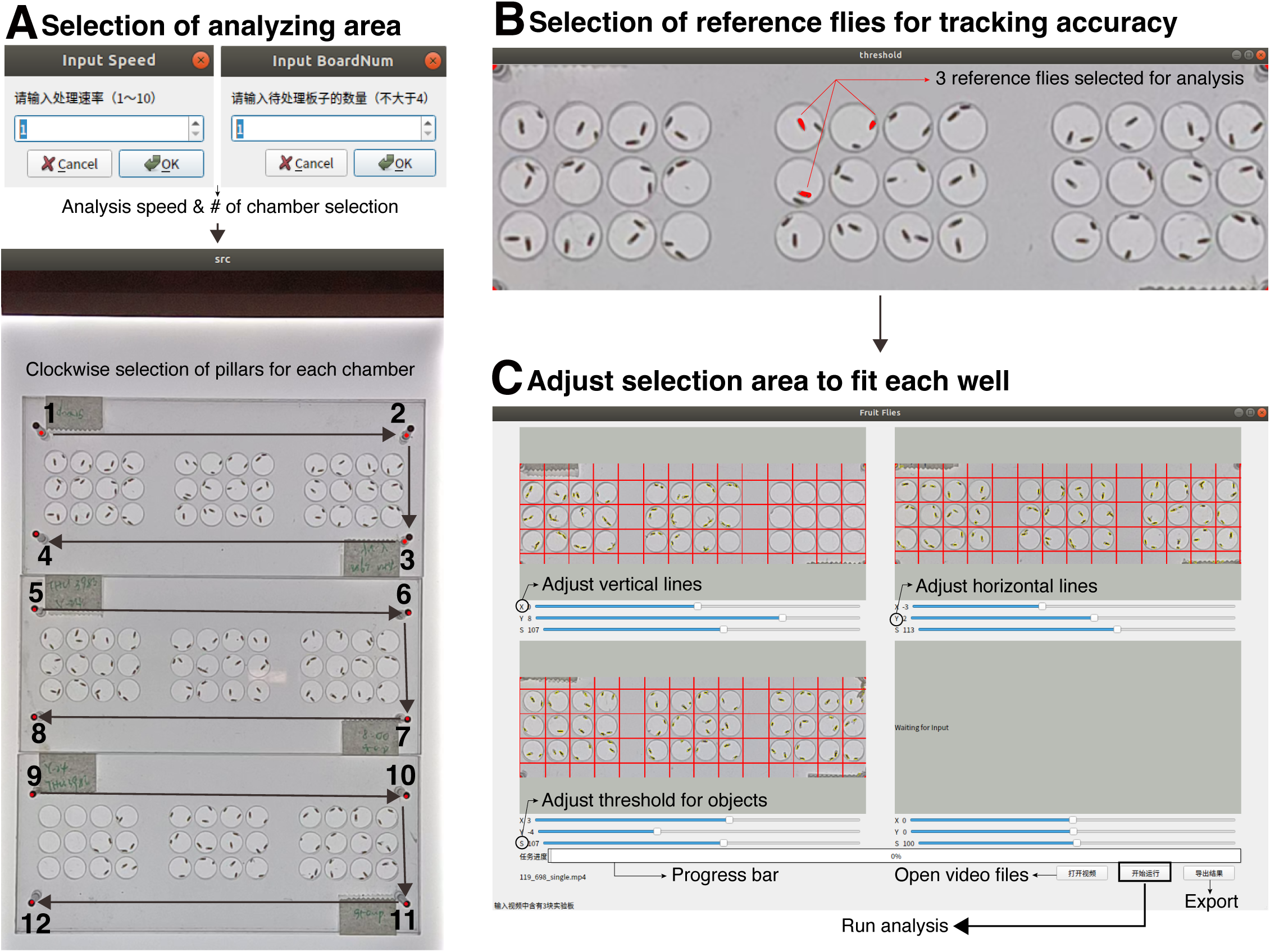
Usage of Linux version of DrosoMating. **(A)** Typing input speed and input board numbers before selecting analyzing area (upper). Schematic representation of the region selection process (lower). Columns should be selected in a clockwise direction starting from the top left corner. **(B)** Selection of reference flies for tracking accuracy. Any three reference flies were chosen for the analysis. **(C)** The DrosoMating home interface. In brief, “Open videos” to import the recorded mating video. “Run analysis” to initiate the analysis process. The progress bar indicates the status of the analysis. Export raw data by clicking “Export”.

**Figure S3.**
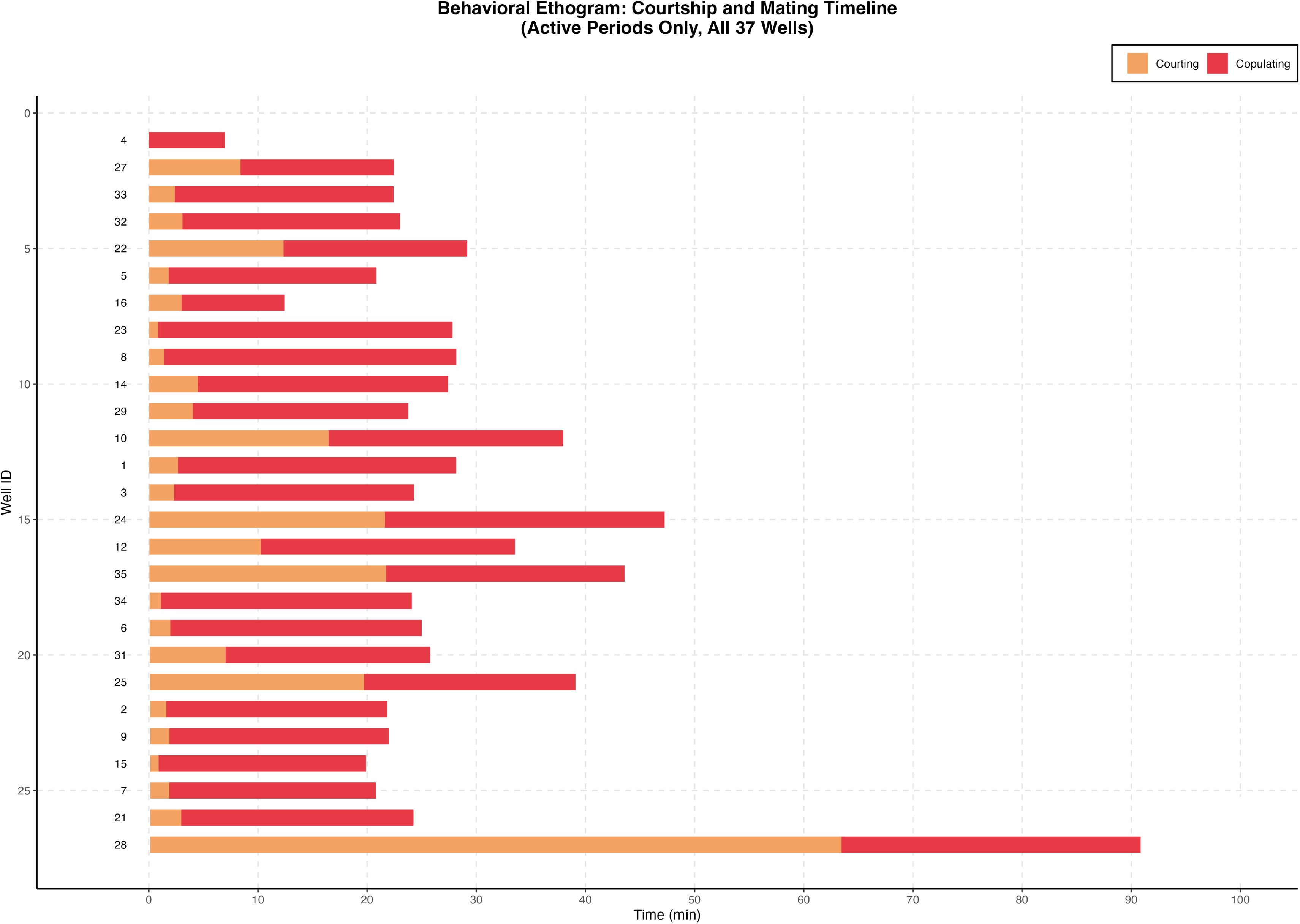
Behavioral ethogram of courtship and copulation dynamics across individual wells. Horizontal bars depict the temporal progression of male mating behaviors. Orange indicates courtship, red indicates copulation, and the x□axis represents time in minutes. Each row corresponds to a single well labeled with well ID on the left. Only wells with successful copulation are shown, sorted by courtship onset.

**Figure S4.**
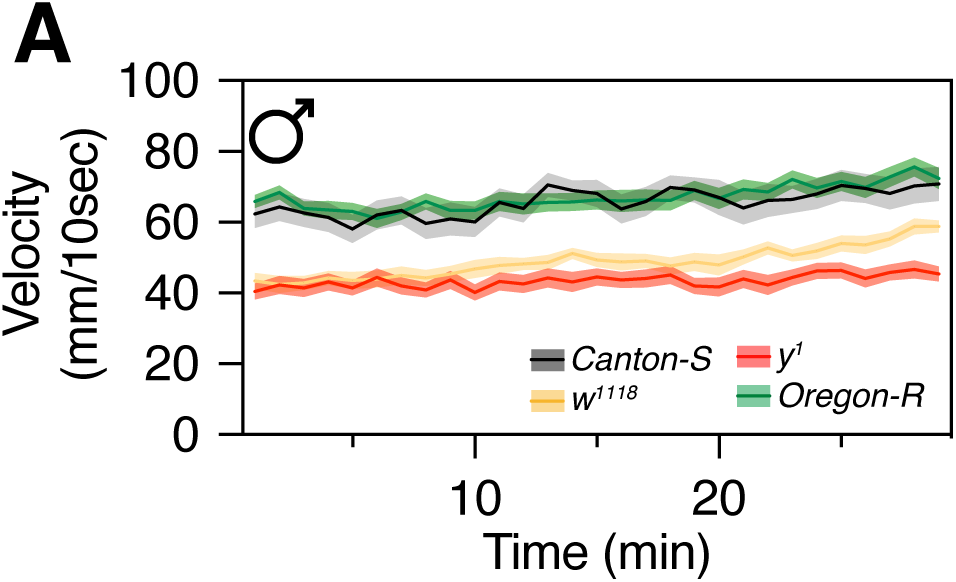
Basal locomotor activity of male flies from different strains. (A) Average locomotor velocity of single-housed male flies from Canton-S, Oregon-R, w^1118^, and y^1^ strains measured in the absence of females. Velocity was quantified over time and plotted as mm per 10 s. Compared with Canton-S and Oregon-R controls, w^1118^ and y^1^ males exhibited reduced basal locomotor activity, indicating that general motor activity contributes to strain-dependent behavioral differences observed in mating assays.

## REFERENCES

Barwell T, Raina S, Seroude L. 2021. Versatile method to measure locomotion in adult Drosophila1. Genome 64:139–145. doi:10.1139/gen-2020-0044

Benton R. 2015. Neural Circuits: Male Mating Motifs. Neuron 87:912–914. doi:10.1016/j.neuron.2015.08.017

Berg S, Kutra D, Kroeger T, Straehle CN, Kausler BX, Haubold C, Schiegg M, Ales J, Beier T, Rudy M, Eren K, Cervantes JI, Xu B, Beuttenmueller F, Wolny A, Zhang C, Koethe U, Hamprecht FA, Kreshuk A. 2019. ilastik: interactive machine learning for (bio)image analysis. Nat Methods 16:1226–1232. doi:10.1038/s41592-019-0582-9

Branson K, Robie AA, Bender J, Perona P, Dickinson MH. 2009. High-throughput ethomics in large groups of Drosophila. Nat Methods 6:451–457. doi:10.1038/nmeth.1328

Bretman A, Fricke C, Chapman T. 2009. Plastic responses of male Drosophila melanogaster to the level of sperm competition increase male reproductive fitness. Proc Royal Soc B Biological Sci 276:1705–1711. doi:10.1098/rspb.2008.1878

Bretman A, Westmancoat JD, Gage MJG, Chapman T. 2011. Males Use Multiple, Redundant Cues to Detect Mating Rivals. Curr Biol 21:617–622. doi:10.1016/j.cub.2011.03.008

Chen C-H, Lin Y-C, Wang S-H, Kuo T-H, Tsai H-Y. 2024a. An automatic system for recognizing fly courtship patterns via an image processing method. Behav Brain Funct 20:5. doi:10.1186/s12993-024-00231-4

Claridge-Chang A, Assam PN. 2016. Estimation statistics should replace significance testing. Nat Methods 13:108–109. doi:10.1038/nmeth.3729

Dankert H, Wang L, Hoopfer ED, Anderson DJ, Perona P. 2009b. Automated monitoring and analysis of social behavior in Drosophila. Nat Methods 6:297–303. doi:10.1038/nmeth.1310

Dauwalder B. 2008. Systems Behavior: Of Male Courtship, the Nervous System and Beyond in Drosophila. Curr Genom 9:517–524. doi:10.2174/138920208786847980

Drapeau MD, Cyran SA, Viering MM, Geyer PK, Long AD. 2006. A cis-regulatory Sequence Within the yellow Locus of Drosophila melanogaster Required for Normal Male Mating Success. Genetics 172:1009–1030. doi: 10.1534/genetics.105.045666

Dukas R. 2020. Natural history of social and sexual behavior in fruit flies. Sci Rep 10:21932. doi:10.1038/s41598-020-79075-7

Eastwood L, Burnet B. 1977a. Courtship latency in maleDrosophila melanogaster. Behav Genet 7:359–372. doi:10.1007/bf01077449

Ejima A, Griffith pring Harb Protoc 2007:pdb.prot4847. doi:10.1101/pdb.prot4LC. 2007. Measurement of Courtship Behavior in Drosophila melanogaster. Cold S847

Eyjolfsdottir E, Branson S, Burgos-Artizzu XP, Hoopfer ED, Schor J, Anderson DJ, Perona P. 2014. Detecting Social Actions of Fruit Flies. Lect Notes Comput Sci 8692:772--787. doi:10.1007/978-3-319-10593-2_50

Gal A, Saragosti J, Kronauer DJ. 2020. anTraX, a software package for high-throughput video tracking of color-tagged insects. eLife 9:e58145. doi:10.7554/elife.58145

Gil-Martí B, Barredo CG, Pina-Flores S, Poza-Rodriguez A, Treves G, Rodriguez-Navas C, Camacho L, Pérez-Serna A, Jimenez I, Brazales L, Fernandez J, Martin FA. 2023a. A simplified courtship conditioning protocol to test learning and memory in Drosophila. Star Protoc 4:101572. doi:10.1016/j.xpro.2022.101572

Greenspan RJ, Ferveur J-F. 2000a. COURTSHIP IN DROSOPHILA. Annu Rev Genet 34:205–232. doi:10.1146/annurev.genet.34.1.205

Griffith LC, Ejima A. 2009a. Courtship learning in Drosophila melanogaster: Diverse plasticity of a reproductive behavior. Learn Mem 16:743–750. doi:10.1101/lm.956309

Hall JC. 1994. The Mating of a Fly. Science 264:1702–1714. doi:10.1126/science.8209251 https://github.com/hcls-kimlab/DrosoMating. n.d.

Huang Y, Kwan A, Kim WJ. 2024. Y chromosome genes interplay with interval timing in regulating mating duration of male Drosophila melanogaster. Gene Rep 101999. doi:10.1016/j.genrep.2024.101999

Iyengar A, Imoehl J, Ueda A, Nirschl J, Wu C-F. 2012. Automated Quantification of Locomotion, Social Interaction, and Mate Preference in Drosophila Mutants. J Neurogenet 26:306–316. doi:10.3109/01677063.2012.729626

Kabra M, Robie AA, Rivera-Alba M, Branson S, Branson K. 2013. JAABA: interactive machine learning for automatic annotation of animal behavior. Nat Methods 10:64–67. doi:10.1038/nmeth.2281

Kamyshev NG, Iliadi KG, Bragina JV. 1999. Drosophila Conditioned Courtship: Two Ways of Testing Memory. Learn Mem 6:1–20. doi:10.1101/lm.6.1.1

Kang K, Panzano VC, Chang EC, Ni L, Dainis AM, Jenkins AM, Regna K, Muskavitch MAT, Garrity PA. 2012. Modulation of TRPA1 thermal sensitivity enables sensory discrimination in Drosophila. Nature 481:76–80. doi:10.1038/nature10715

Kim WJ, Jan LY, Jan YN. 2013a. A PDF/NPF Neuropeptide Signaling Circuitry of Male Drosophila melanogaster Controls Rival-Induced Prolonged Mating. Neuron 80:1190–1205. doi:10.1016/j.neuron.2013.09.034

Kim WJ, Jan LY, Jan YN. 2012a. Contribution of visual and circadian neural circuits to memory for prolonged mating induced by rivals. Nat Neurosci 15:876–883. doi:10.1038/nn.3104

Kim WJ, Lee SG, Schweizer J, Auge A-C, Jan LY, Jan YN. 2016. Sexually experienced male Drosophila melanogaster uses gustatory-to-neuropeptide integrative circuits to reduce time investment for mating. Biorxiv 088724. doi:10.1101/088724

Kim WJ, Song Y, Zhang T, Zhang X, Ryu TH, Wong KC, Wu Z, Wei Y, Schweizer J, Nguyen K-NH, kwan A, Yu K. 2024. Peptidergic neurons with extensive branching orchestrate the internal states and energy balance of male Drosophila melanogaster. bioRxiv 2024.06.04.597277. doi:10.1101/2024.06.04.597277

Kim Y-K. 2009. Handbook of Behavior Genetics 317–330. doi:10.1007/978-0-387-76727-7_22

Kitamoto T. 2001. Conditional modification of behavior in Drosophila by targeted expression of a temperature-sensitive shibire allele in defined neurons. J Neurobiol 47:81–92. doi:10.1002/neu.1018

Krstic D, Boll W, Noll M. 2013. Influence of the White Locus on the Courtship Behavior of Drosophila Males. PLoS ONE 8:e77904. doi:10.1371/journal.pone.0077904

Lee SG, Kang C, Saad B, Nguyen K-NH, Guerra-Phalen A, Bui D, Abbas A-H, Trinh B, Malik A, Zeghal M, Auge A-C, Islam ME, Wong K, Stern T, Lebedev E, Sun D, Miao H, Wu Z, Sherratt TN, Kim WJ. 2022. Multisensory inputs control the regulation of time investment for mating by sexual experience in male Drosophila melanogaster. Biorxiv 2022.09.23.509131. doi:10.1101/2022.09.23.509131

Lee SG, Sun D, Miao H, Wu Z, Kang C, Saad B, Nguyen K-NH, Guerra-Phalen A, Bui D, Abbas A-H, Trinh B, Malik A, Zeghal M, Auge A-C, Islam ME, Wong K, Stern T, Lebedev E, Sherratt TN, Kim WJ. 2023a. Taste and pheromonal inputs govern the regulation of time investment for mating by sexual experience in male Drosophila melanogaster. PLOS Genet 19:e1010753. doi:10.1371/journal.pgen.1010753

Levin LR, Han P-L, Hwang PM, Feinstein PG, Davis RL, Reed RR. 1992. The Drosophila learning and memory gene rutabaga encodes a Ca2+calmodulin-responsive adenylyl cyclase. Cell 68:479–489. doi:10.1016/0092-8674(92)90185-f

Neckameyer WS, Bhatt P. 2016. Drosophila, Methods and Protocols. Methods Mol Biol 1478:303–320. doi:10.1007/978-1-4939-6371-3_19

Pavlou HJ, Goodwin SF. 2013. Courtship behavior in Drosophila melanogaster: towards a ‘courtship connectome.’ Curr Opin Neurobiol 23:76–83. doi:10.1016/j.conb.2012.09.002

Pérez-Escudero A, Vicente-Page J, Hinz RC, Arganda S, Polavieja GG de. 2014. idTracker: tracking individuals in a group by automatic identification of unmarked animals. Nat Methods 11:743–748. doi:10.1038/nmeth.2994

Qu S, Zhu Q, Zhou H, Gao Y, Wei Y, Ma Y, Wang Z, Sun X, Zhang Lei, Yang Q, Kong L, Zhang Li. 2022. EasyFlyTracker: A Simple Video Tracking Python Package for Analyzing Adult Drosophila Locomotor and Sleep Activity to Facilitate Revealing the Effect of Psychiatric Drugs. Front Behav Neurosci 15:809665. doi:10.3389/fnbeh.2021.809665

Reza MdA, Mhatre SD, Morrison JC, Utreja S, Saunders AJ, Breen DE, Marenda DR. 2013. Automated analysis of courtship suppression learning and memory in Drosophila melanogaster. Fly 7:105–111. doi:10.4161/fly.24110

Risse B, Berh D, Otto N, Klämbt C, Jiang X. 2017. FIMTrack: An open source tracking and locomotion analysis software for small animals. PLoS Comput Biol 13:e1005530. doi:10.1371/journal.pcbi.1005530

Risse B, Otto N, Berh D, Jiang X, Klämbt C. 2014. FIM Imaging and FIMtrack: Two New Tools Allowing High-throughput and Cost Effective Locomotion Analysis. J Vis Exp. doi:10.3791/52207-v

Rodriguez A, Zhang H, Klaminder J, Brodin T, Andersson PL, Andersson M. 2017. ToxTrac_: A fast and robust software for tracking organisms. Methods Ecol Evol 9:460–464. doi:10.1111/2041-210x.12874

Simon AF, Chou M-T, Salazar ED, Nicholson T, Saini N, Metchev S, Krantz DE. 2011. A simple assay to study social behavior in Drosophila: measurement of social space within a group. Genes, brain, Behav 11:243–52. doi:10.1111/j.1601-183x.2011.00740.x

Singh A, Singh BN. 2014. Mating latency, duration of copulation and fertility in four species of the Drosophila bipectinata complex. Indian J Exp Biol 52:175–80.

Singh BN, Singh A. 2016. The genetics of sexual behavior in Drosophila. Adv Genom Genet 6:1–9. doi:10.2147/agg.s58525

Sokolowski MB. 2001. Drosophila: Genetics meets behaviour. Nat Rev Genet 2:879–890. doi:10.1038/35098592

Stern U, Zhu EY, He R, Yang C-H. 2015. Long-duration animal tracking in difficult lighting conditions. Sci Rep 5:10432. doi:10.1038/srep10432

Sun D, Zhang X, Miao H, Zhang T, Kim WJ. 2024. Molecular and Neural Circuit Mechanisms Underlying Sexual Experience-dependent Long-Term Memory in Drosophila. bioRxiv 2024.09.28.615582. doi:10.1101/2024.09.28.615582

Sun Y, Zhang X, Wu Z, Li W, Kim WJ. 2024. Genetic Screening Reveals Cone Cell-Specific Factors as Common Genetic Targets Modulating Rival-Induced Prolonged Mating in male Drosophila melanogaster. G3: Genes, Genomes, Genet jkae255. doi:10.1093/g3journal/jkae255

Vosshall LB. 2007. Into the mind of a fly. Nature 450:193–197. doi:10.1038/nature06335

Wolf R, Wittig T, Liu L, Wustmann G, Eyding D, Heisenberg M. 1998. Drosophila Mushroom Bodies Are Dispensable for Visual, Tactile, and Motor Learning. Learn Mem 5:166–178. doi:10.1101/lm.5.1.166

Wong K, Schweizer J, Nguyen K-NH, Atieh S, Kim WJ. 2019. Neuropeptide relay between SIFa signaling controls the experience-dependent mating duration of male Drosophila. Biorxiv 819045. doi:10.1101/819045

Yamamoto D, Koganezawa M. 2013. Genes and circuits of courtship behaviour in Drosophila males. Nat Rev Neurosci 14:681–692. doi:10.1038/nrn3567

Yamanaka O, Takeuchi R. 2018. UMATracker: an intuitive image-based tracking platform. J Exp Biol 221:jeb182469. doi:10.1242/jeb.182469

Yapici N, Kim Y-J, Ribeiro C, Dickson BJ. 2008. A receptor that mediates the post-mating switch in Drosophila reproductive behaviour. Nature 451:33–37. doi:10.1038/nature06483

Zhang T, Wu Z, Song Y, Li W, Sun Y, Zhang X, Wong K, Schweizer J, Nguyen K-NH, Kwan A, Kim WJ. 2024a. Long-range neuropeptide relay as a central-peripheral communication mechanism for the context-dependent modulation of interval timing behaviors. bioRxiv 2024.06.03.597273. doi:10.1101/2024.06.03.597273

Zhang T, Zhang X, Sun D, Kim WJ. 2024b. Exploring the Asymmetric Body’s Influence on Interval Timing Behaviors of Drosophila melanogaster. Behav Genet 1–10. doi:10.1007/s10519-024-10193-y

Zhang X, Miao H, Kang D, Sun D, Kim WJ. 2024. Male-specific sNPF peptidergic circuits control energy balance for mating duration through neuron-glia interactions. bioRxiv 2024.10.17.618859. doi:10.1101/2024.10.17.618859

